# IL-6 underlies microenvironment immunosuppression and resistance to therapy in glioblastoma

**DOI:** 10.1101/2025.03.12.642800

**Authors:** Jacob S. Young, Nam Woo Cho, Calixto-Hope G. Lucas, Hinda Najem, Kanish Mirchia, William C. Chen, Kyounghee Seo, Naomi Zakimi, Vikas Daggubati, Tim Casey-Clyde, Minh P. Nguyen, Arya Chen, Joanna J. Phillips, Tomoko Ozawa, Manish K. Aghi, Jennie W. Taylor, Joseph L. DeRisi, Aparna Bhaduri, Mitchel S. Berger, Amy B. Heimberger, Nicholas Butowski, Matthew H. Spitzer, David R. Raleigh

## Abstract

The glioblastoma tumor immune microenvironment (TIME) is an immunosuppressive barrier to therapy that encumbers glioblastoma responses to immune checkpoint inhibition (ICI). Immunosuppressive cytokines, pro-tumor myeloid cells, and exhausted T-cells are hallmarks of the glioblastoma TIME. Here we integrate spatial and single-cell analyses of patient-matched human glioblastoma samples before and after ICI with genetic, immunologic, single-cell, and pharmacologic studies in preclinical models to reveal that interleukin-6 (IL-6) inhibition reprograms the glioblastoma TIME to sensitize mouse glioblastoma to ICI and radiotherapy. Rare human glioblastoma patients who achieve clinical responses to ICI have lower pre-treatment IL-6 levels compared to glioblastomas who do not respond to ICI. Immune stimulatory gene therapy suppresses IL-6 tumor levels in preclinical murine models of glioblastoma. Furthermore, survival was longer in *Il-6* knockout mice with orthotopic SB28 glioblastoma relative to wild-type mice. IL-6 blockade with a neutralizing antibody transiently sensitizes mouse glioblastoma to anti-PD-1 by increasing MHCII+ monocytes, CD103+ migratory dendritic cells (DCs), CD11b+ conventional DCs, and effector CD8+ T cells, and decreasing immunosuppressive Tregs. To translate these findings to a combination treatment strategy for recurrent glioblastoma patients, we show that IL-6 blockade plus ICI durably sensitizes mouse glioblastoma to high-dose radiotherapy.

## Introduction

Glioblastoma is the most aggressive central nervous system (CNS) malignancy in adults and is associated with poor survival despite aggressive treatment with surgery, ionizing radiation, and chemotherapy^1^. Immune checkpoint inhibition (ICI) has revolutionized treatments for solid and hematological malignancies, but durable responses to ICI are lacking in glioblastoma^2–4^. Glioblastomas are considered immunologically “cold” tumors that have an immunosuppressive tumor immune microenvironment (TIME) mostly comprised of pro-tumor macrophages and myeloid cells^5–10^. Moreover, daily low-dose radiotherapy, chemotherapy, and corticosteroids, which form the backbone of adjuvant treatment for glioblastoma, contribute to immunosuppression systemically and in the TIME^11^, and attempts to overcome this barrier with combination treatments that are effective in other cancers have not increased survival from glioblastoma^12^.

The interleukin-6 (IL-6) signaling axis is classically considered pro-inflammatory and drives autoimmunity^13^. Paradoxically, IL-6 in the glioblastoma TIME can stimulate the growth of glioma stem cells and increases immunosuppressive cytokines that activate pro-tumor macrophages^14–18^. IL-6 signaling has also been implicated in the development of resistance to treatment, suggesting that IL-6 blockade may improve ICI responses in cancers like glioblastoma^19^. In mouse models of non-CNS cancer, IL-6 blockade reduces the number of macrophages, myeloid cells, and Th17 cells and increases the number of effector T cells in the TIME, leading to therapeutic synergy when combined with ICI^20^. IL-6 signaling in human glioblastoma patients after treatment with ICI has not been investigated, and the impact of IL-6 blockade on the glioblastoma TIME and its potential synergy with ICI or other adjuvant therapies are incompletely understood.

Here we investigate glioblastoma ICI responses using single-cell and spatial analyses of human samples and preclinical mouse glioblastoma models after treatment with experimental or standard-of-care therapeutics. Our results reveal that endogenous IL-6 and downstream IL-6 effectors that support cancer stemness^21^ are associated with poor clinical responses to ICI, and that combined IL-6 blockade and ICI in mice induces CD8+ T-cell expansion while suppressing CD4+ regulatory T cell (Treg) infiltration to reprogram the TIME and improve overall survival. To translate these findings to a therapeutically relevant combination treatment for patients, we show that IL-6 blockade plus ICI durably prolongs survival in preclinical models after ablative radiotherapy. These data suggest that IL-6 and the IL-6 signaling cascade contribute to ICI resistance and shed light on a combination treatment strategy for the most aggressive CNS malignancy in adults.

## Results

### Glioblastomas that respond to immune checkpoint inhibition are enriched in T cells after treatment

To define mechanisms underlying glioblastoma responses to ICI, spatial protein profiling was performed on 14 patient-matched glioblastoma samples (IDH-wildtype, CNS WHO grade 4) from 7 patients who underwent surgery before and after treatment with ICI at the University of California San Francisco (Supplementary Table 1). Magnetic resonance imaging (MRI) studies, histopathological reports, next-generation sequencing of tumor tissue, and medical records were reviewed to distinguish ICI responders (n=3) from ICI non-responders (n=4) (Fig. 1a, Supplementary Table 2, Methods). As previously reported, ICI responders were defined (1) by the presence of robust reactive immune infiltrate with scant or no tumor cells after ICI, or (2) tumor size stability or decreasing tumor size over at least 6 months from initiation of ICI on MRI^22,23^. Spatial protein profiling of 200μm regions-of-interest corresponding to viable tumor was performed on 6mm cores from formalin-fixed paraffin-embedded (FFPE) tumor tissue blocks for all samples (Fig. 1b, Extended Data Fig. 1a, b). Laser microdissection and polymerase chain reaction sequencing were used to quantify the binding of 78 antibodies that were conjugated to unique oligonucleotide barcodes across an average of 6 spatial regions/sample (n=84 total regions) (Supplementary Table 3), which were selected to avoid interrogation of tumor necrosis.

**Fig. 1.**
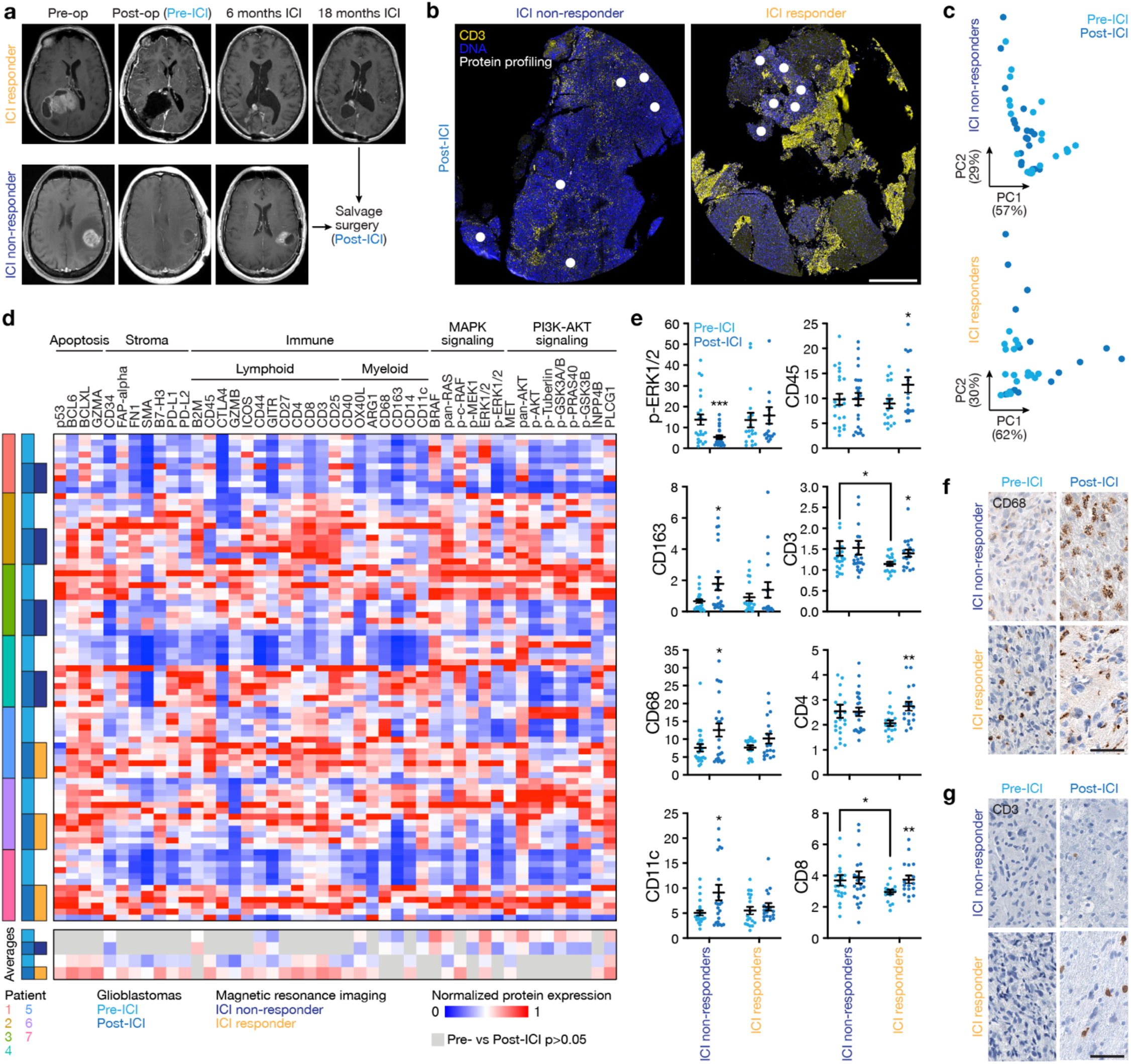
Glioblastomas that respond to immune checkpoint inhibition are enriched in intratumor T cells after treatment. **a**, 7 patients with 14 match-paired glioblastoma samples that were resected before or after treatment with ICI. Responders (top) and non-responders (bottom) were distinguished using radiographic and pathologic criteria (Methods). **b**, CD3 and DAPI IF sections from an ICI non-responder (left) or an ICI responder (right) showing regions of spatial protein profiling using the NanoString GeoMx Digital Spatial Profiler in viable tumor tissue (white) adjacent to regions of intratumor necrosis. Representative of n=7 patients. Scale bar, 1mm. **c**, Principal component (PC) analysis of expression from spatial protein profiling of ICI non-responders (top, n=4 patients) or ICI responders (bottom, n=3 patients). **d**, Heatmaps of differentially expressed spatial proteins revealing significant inter- and intratumor heterogeneity between ICI responders and non-responders. Top heatmap shows results from 84 spatial regions across 14 matched-paired samples from 7 patients. Bottom heatmap shows significantly different results (Student’s t tests, p≤0.05) for ICI responders or ICI non-responders when comparing pre-ICI vs post-ICI samples. **e**, Spatial protein expression of select immune or MAPK signaling markers showing post-ICI myeloid infiltration in non-responders and post-ICI lymphoid infiltration in responders. Student’s t tests, *p≤0.05, **p≤0.01. **f**, CD68 IHC sections from pre-ICI (left) or post-ICI (right) samples from ICI non-responders (top) or ICI responders (bottom). Representative of n=7 patients. **g**, CD3 IHC sections from pre-ICI (left) or post-ICI (right) samples from ICI non-responders (top) or responders (bottom). Scale bar, 100µm. Representative of n=7 patients.

Spatial protein profiling demonstrated key immunologic signaling and cellular differences between pre- and post-treatment glioblastoma samples from ICI responders versus ICI non-responders (Fig. 1c-e). Principle component analysis of spatial protein profiling data suggested divergent biochemical factors post-ICI compared to pre-ICI in ICI responders compared to ICI non-responders (Fig. 1c). Proteins associated with apoptosis (p53, BCL6) and lymphoid cell infiltration and activation (CD3, CD4, CD8, CD25, CD40) were enriched in ICI responders after ICI treatment, whereas proteins associated with myeloid cell infiltration (CD163, CD68, CD11c) were enriched in ICI non-responders after ICI treatment (Fig. 1d, e, Supplementary Table 3). Downstream markers of IL-6 signaling such as MAPK activation (p-ERK1/2) and PI3K-AKT activation (p-AKT)^24–26^ were enriched in ICI non-responders before ICI treatment (Fig. 1d, e). Immunohistochemical staining for CD68 and CD3 on tissue sections validated changes in myeloid and lymphoid cells that had been identified using spatial protein profiling (Fig. 1f, g). These data show that glioblastoma ICI responders achieve lymphocytic immune cell infiltration, while glioblastoma ICI non-responders are infiltrated with myeloid cells after treatment and are enriched in MAPK and PI3K-AKT signaling before treatment, both of which are modulated by IL-6.

### Glioblastomas that do not respond to immune checkpoint inhibition are enriched in immunosuppressive myeloid cells and cancer stem cells after treatment

Spatial transcriptomic profiling of 200μm regions from viable tumor tissue on FFPE sections demonstrated gene expression differences between pre- and post-treatment glioblastoma samples from ICI responders versus ICI non-responders (Fig. 2a-e, Supplementary Table 4). Differential expression analysis showed enrichment of stem cell markers (*SOX13, SOX4, PTPRZ1*) and innate immune markers (*IFI44L, IFI27, IFITM1*) in ICI non-responders after treatment (Fig. 2b-e) and suppression of stem cell markers (*SOX4, SOX9, PTPRZ1*) in ICI responders after treatment (Fig. 2b-e), which was confirmed by gene ontology analyses (Fig. 2a, d, Extended Data Fig. 2a). Notably, IL-6 and the IL-6 signaling cascade support glioblastoma tumorigenesis and stemness^21,27^, and we identified that genes involved in the IL-6 signaling pathway, such as *ANXA1*, *LGALS3*, and *EGFR*, were enriched in ICI non-responders pre-ICI (Fig. 2c)^28–30^.

**Fig. 2.**
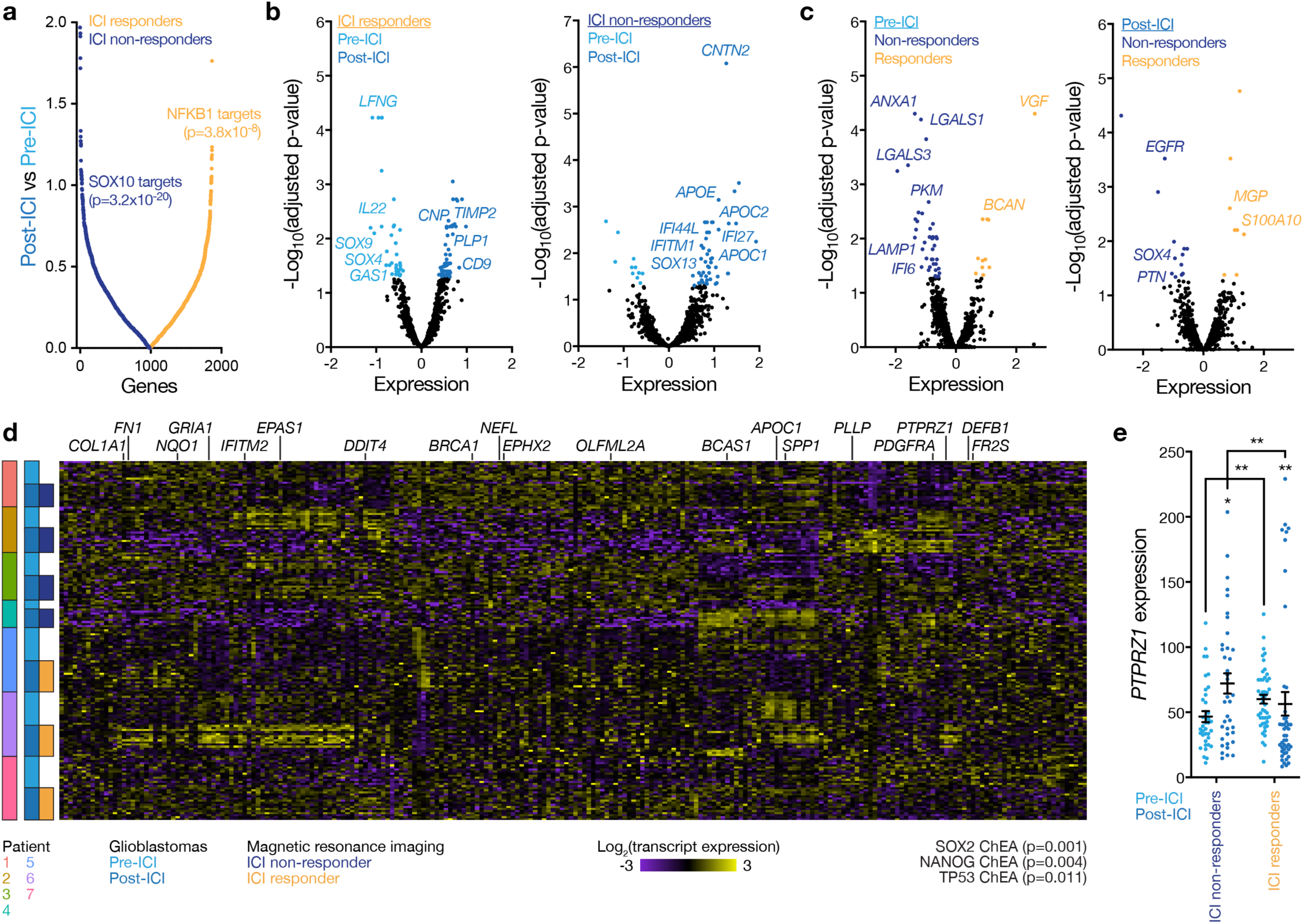
Glioblastomas that do not respond to immune checkpoint inhibition are enriched in intratumor innate immune and cancer stem cell markers after treatment. **a**, Differential spatial transcriptomic expression from 7 patients with 14 match-paired glioblastoma samples that were resected before or after treatment with ICI. Differentially expressed spatial genes in viable tumor tissue adjacent to regions of intratumor necrosis, ranked by magnitude of expression, are shown. ENRICHR gene ontology terms using the top 250 differentially expressed spatial genes in post-ICI vs pre-ICI samples from non-responders (left) or responders (right) are also shown. **b**, Volcano plots showing differential spatial gene expression in pre-ICI vs post-ICI samples from ICI responders (left) or ICI non-responders (right). **c**, Volcano plots showing differential spatial gene expression in ICI non-responders vs ICI responders from pre-ICI (left) or post-ICI (right) samples. **d**, Heatmaps of differentially expressed spatial genes from 173 regions revealing significant inter- and intratumor heterogeneity between ICI responders and ICI non-responders in pre-ICI vs post-ICI samples. ENRICHR gene ontology terms using the top 250 differentially expressed spatial genes across all samples are shown. **e**, Spatial expression of the glioblastoma stem cell marker gene *PTPRZ1* shows enrichment in ICI responders vs ICI non-responders pre-ICI and in post-ICI vs pre-ICI samples from non-responders. *PTPRZ1* was suppressed in ICI responders vs ICI non-responders post-ICI and in post-ICI vs pre-ICI samples from responders. Student’s t tests, *p≤0.05, **p≤0.01.

To further investigate the signaling mechanisms and immunophenotypes in glioblastomas before and after treatment with ICI, we performed multiplexed sequential immunofluorescence (seq-IF) to define cellular phenotypes underlying spatial protein and gene expression programs in ICI responders and non-responders (Fig. 3a, Supplementary Table 5). ICI non-responders were enriched in glioblastoma stem cells marked by SOX2 and PTPRZ1 and in cells marked by the IL-6 effector p-STAT3 in both pre- and post-ICI samples compared to ICI responders (Fig. 3b). ICI non-responders were also enriched in myeloid cells marked by CD68, CD163, and CD11c, as well as microglia marked by P2RY12, in both pre- and post-ICI samples compared to ICI responders (Fig. 3c). Lymphoid cells were scarce in both ICI responders and ICI non-responders compared to glioblastoma stem cell and myeloid cell populations in non-responders using seq-IF (Fig. 3d). In sum, spatial protein and transcriptomic profiling and seq-IF of human glioblastomas before and after treatment with ICI suggest that IL-6 may support glioblastoma tumorigenesis and stemness to drive resistance to therapy through target genes like *ANXA1* and *EGFR*, and pathway effectors such a p-ERK1/2, p-AKT, and p-STAT3. To test this hypothesis, we turned to preclinical models to determine if the immunosuppressive TIME could be reprogrammed to sensitize tumors to ICI.

**Fig. 3.**
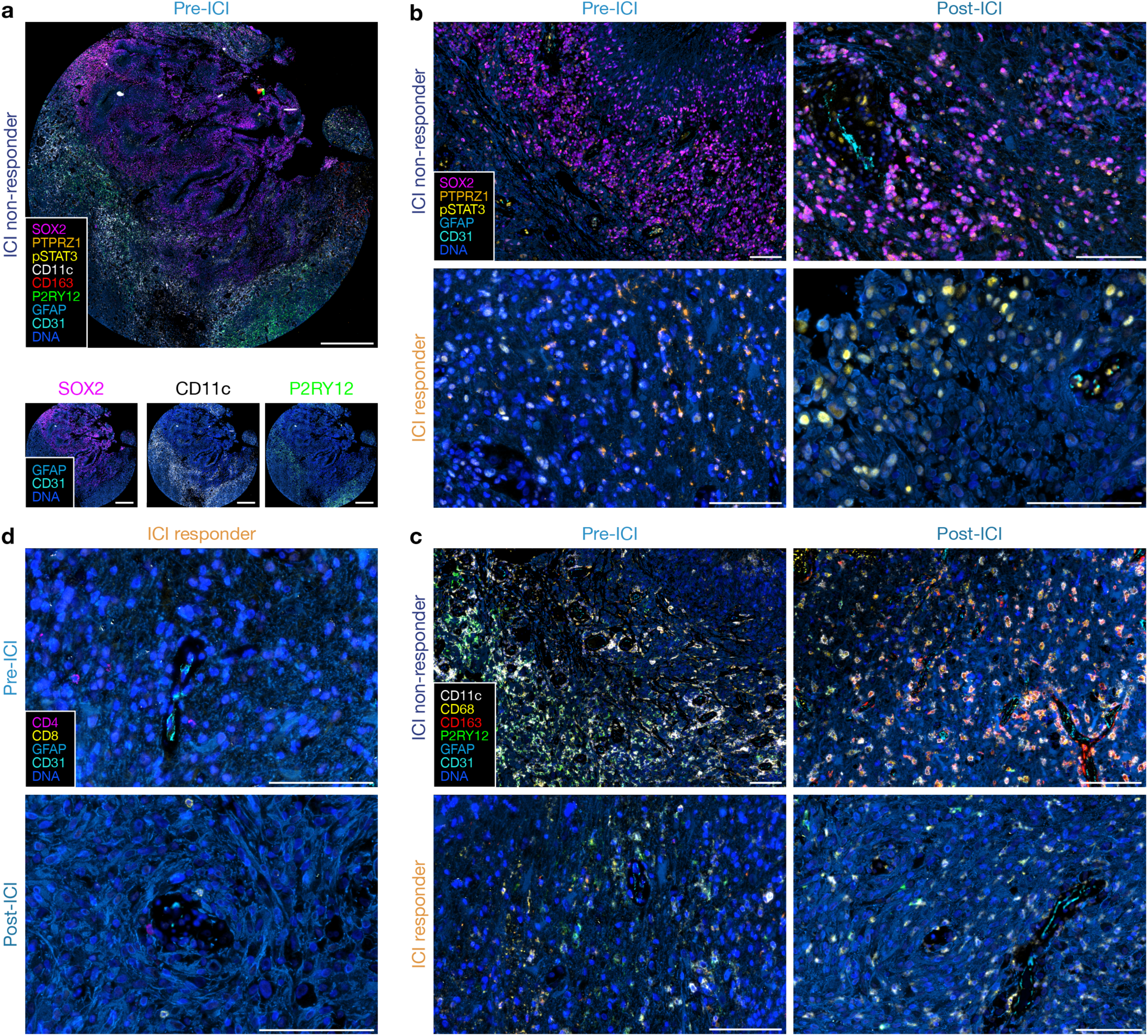
Glioblastomas that do not respond to immune checkpoint inhibition are enriched in immunosuppressive myeloid cells and glioma stem-like cells after treatment. a, Multiplexed sequential immunofluorescence microscopy showing intratumor heterogeneity of signaling mechanisms and cell types in ICI non-responder tissue section. Scale bar, 1mm. **b,** Multiplexed sequential immunofluorescence microscopy on the Lunaphore Comet platform showing dramatic increase in SOX2+, PTPRZ1+, and p-STAT3+ expressing cells in ICI non-responders. Scale bar, 100µm. **c,** Multiplexed sequential immunofluorescence microscopy showing few myeloid cells in ICI responders, particularly CD163+ macrophages and P2RY12+ myeloid cells, compared to ICI non-responders. Scale bar, 100µm. **d,** Multiplexed sequential immunofluorescence microscopy showing rare naïve perivascular lymphocytes in ICI responders both pre- and post-ICI treatment. Scale bar, 100µm. Images are representative of protein expression program differences across matched pairs of pre-ICI and post-ICI glioblastomas.

### Tumor immune microenvironments of mouse glioblastoma are heterogeneous

Immunocompetent intracranial mouse models of glioblastomas can have dramatic differences in TIME composition, all of which may not accurately recapitulate glioblastomas in humans^31,32^. GL261 and SB28 both generate highly cellular tumors, but GL261 have marked lymphoid cell infiltration whereas SB28 is enriched in myeloid cells, similar to glioblastomas in humans (Fig. 4a)^7,33^. Convection-enhanced delivery (CED) of gene therapy vectors encoding immunostimulatory products can provide therapeutic gene coverage of intra- and peri-tumoral regions in preclinical models (Fig. 4b) and in human subjects with glioblastoma^34–36^. To define the TIME of immunocompetent intracranial mouse models of glioblastoma in the context of intratumor gene therapy vector perturbations and human glioblastomas, single-cell mass cytometry by time-of-flight (CyTOF) was performed on 6 human glioblastomas (IDH-wildtype, CNS WHO grade 4) and 15 intracranial GL261 (n=7) or SB28 (n=8) glioblastoma from C57BL/6J mice with (n=7) or without (n=8) intratumor CED of attenuated adenovirus 9 (AAV9) gene therapy vectors that were modified to express GFP (Supplementary Table 1, 6). Principal component analysis of CyTOF data demonstrated that the TIME of SB28 more closely resembled the TIME of human glioblastomas, whereas there was differential enrichment of CD4+ and CD8+ T cells, NK cells, and dendritic cells in GL261 that was not observed in human glioblastomas (Fig. 4c, d). AAV9-GFP gene therapy CED did not alter the TIME of glioblastoma compared to untreated controls (Fig. 4c). Uniform manifold approximation and projection (UMAP) analysis showed that GL261 was enriched in myeloid cells with iNOS and MHCII+ expression suggesting M1-like anti-tumoral activation, whereas SB28 was enriched in CD206+ macrophages, Ly6C^hi^ monocytes, and MHCII-macrophages that promote immunosuppression^37,38^ (Fig. 4e-g, Extended Data Fig. 3, 4). SB28 also had fewer activated cDCs and MHCII+ microglia compared to GL261 (Fig.4e-g). Within SB28, a greater percentage of lymphoid cells were effector CD8+ T-cells and Ly6C+ activated CD8+ T-cells compared to CD4+ T-cells in the GL261 tumors (Fig. 4e-g). These data suggest that the TIME of SB28 but not GL261 tumors closely resembles the TIME of human glioblastoma ICI non-responders, which comprise most glioblastomas in humans.

**Fig. 4.**
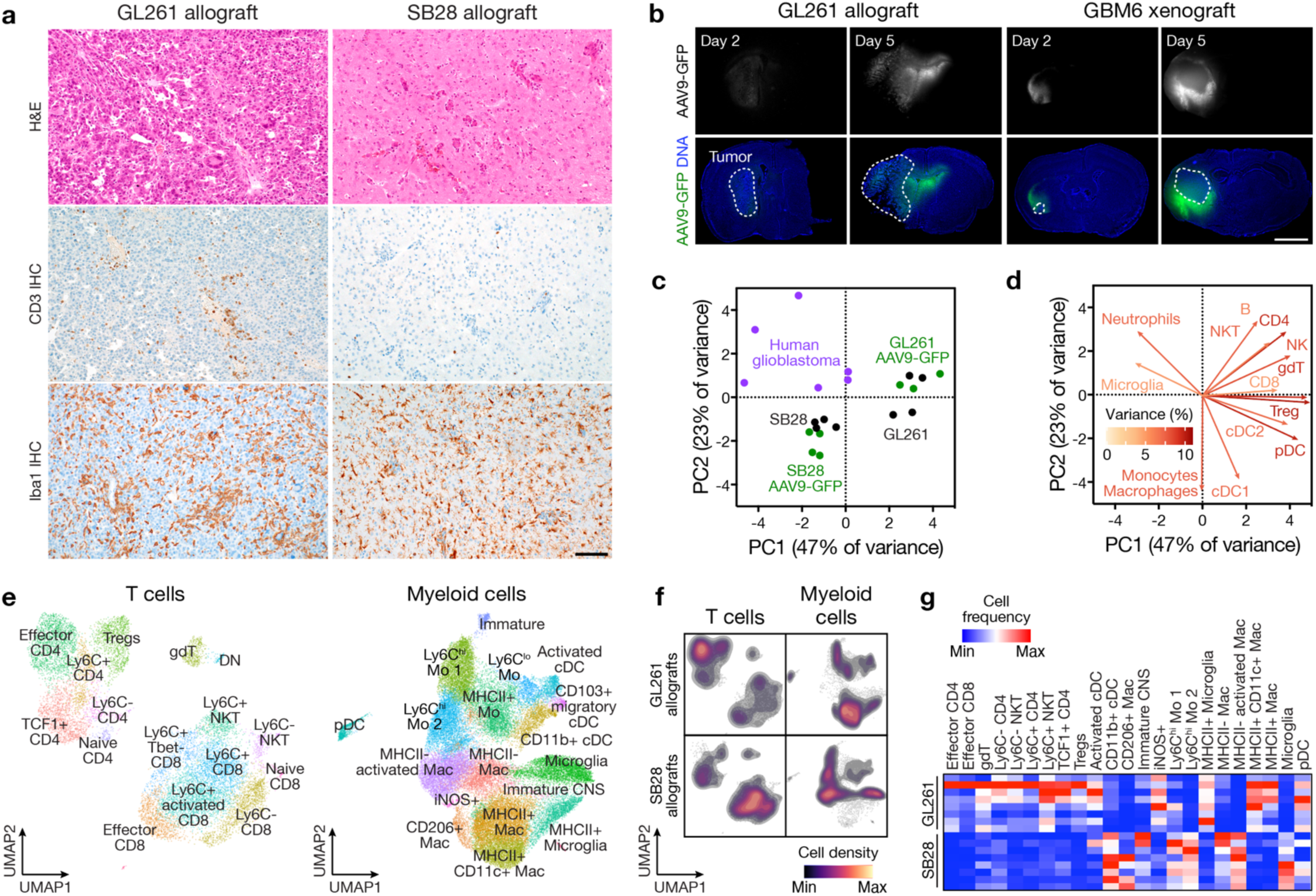
The immune microenvironment of immunocompetent intracranial glioblastoma allografts. **a**, H&E (top), CD3 IHC (middle), and Iba1 IHC (bottom) sections from GL261 (left) or SB28 (right) intracranial glioblastoma allografts in immunocompetent C57BL/6J mice. Representative of n=3 mice/model. Scale bar, 100µm. **b**, IF sections showing intra- and peri-tumor intracranial GFP 2 or 5 days after CED of AAV9-GFP into GL261 allografts (left) or human GBM6 xenografts (right). Representative of n=3 mice/model. Nuclei are marked with DAPI. Scale bar, 1mm. **c**, Principal component (PC) analysis of CD45+ immune cell subset frequencies determined by CyTOF of newly diagnosed human glioblastomas (n=6 patients, 286,370 cells) compared to GL261 (n= 7 mice, 460,176 cells) or SB28 (n=8 mice, 712,187 cells) intracranial glioblastoma allografts after control (n=4 mice/model) or AAV9-GFP (n=3-4 mice/model) treatment revealing the immune microenvironment of human and SB28 glioblastomas are similar. **d**, Variable loadings plot for PC analysis in **c** showing the top 13 immune cell types contributing to variance for PC1 and PC2. Human and SB28 glioblastomas have fewer T cells than GL261 glioblastomas. **e**, UMAP and Phenograph unsupervised clustering analysis of the immune microenvironment in GL261 and SB28 control or AAV9-GFP intracranial glioblastoma allografts for T cells (n=8 mice per model, 12,544 cells, left) or myeloid cells (n=8 mice per model, 40,000 cells, right). **f**, UMAP density plots from analysis in **e** for T cells (left) or myeloid cells (right) in GL261 (top) or SB28 (bottom) intracranial glioblastoma allografts. **g**, Heatmap of T cell or myeloid cell subset frequencies from **e** for individual mice, showing T cell depletion in SB28 glioblastomas. All cell types with Student’s t test p≤0.05 for comparison between models are shown.

### Intratumor cytokine gene therapy reprograms the glioblastoma immune microenvironment

In contrast to human glioblastoma, the TIME of other types of intracranial tumors such as schwannoma are enriched in pro-inflammatory responses including CD4+ and CD8+ effector and memory T-cells and classic dendritic cells. These differences are attributed to tumor-elaborated immunostimulatory cytokines and chemokines such as CXCL2, CXCL3, CCL2, APOA1, IL1B, and CCL4^39,40^. CED of these recombinant cytokines in immunocompetent glioblastoma C57BL/6J mice nominated APOA1, IL-1B, and CCL4 as candidates that could increase immune cell infiltration and necrosis in the intracranial TIME (Fig. 5a). To durably express these cytokines in preclinical glioblastoma models, AAV9 gene therapy vectors encoding *Apoa1, Il1b,* or *Ccl4* were delivered using CED to intracranial SB28 or GL261 in C57BL/6J mice, and the TIME responses analyzed using CyTOF, multiplexed cytokine assays, and histology (Fig. 5b). Intratumor *Apoa1* or *Il1b* gene therapies modestly improved survival in the preclinical SB28 model without apparent toxicity, but there was no survival benefit in the GL261 model (Fig. 5c, Extended Data Fig. 5a). Tumor CyTOF and histology showed T-cell and microglial enrichment in SB28 but not GL261 after *Apoa1* or *Il1b* gene therapy treatments (Fig. 5d, Extended Data Fig. 5b). Multiplexed cytokine assays showed that intratumor gene therapies suppressed IL-6, LIF (an IL-6 cytokine family member), CXCL10, CCL2, and VEGF in SB28 but not in GL261 (Fig. 5e, Extended Data Fig. 5c). Considering the data from preclinical models (Fig. 4, 5) in the context of data from human glioblastomas before and after treatment with ICI (Fig. 1-3), we hypothesized that IL-6 and the IL-6 signaling cascade may drive glioblastoma tumorigenesis and stemness, and that IL-6 blockade may sensitize glioblastomas to ICI.

**Fig. 5.**
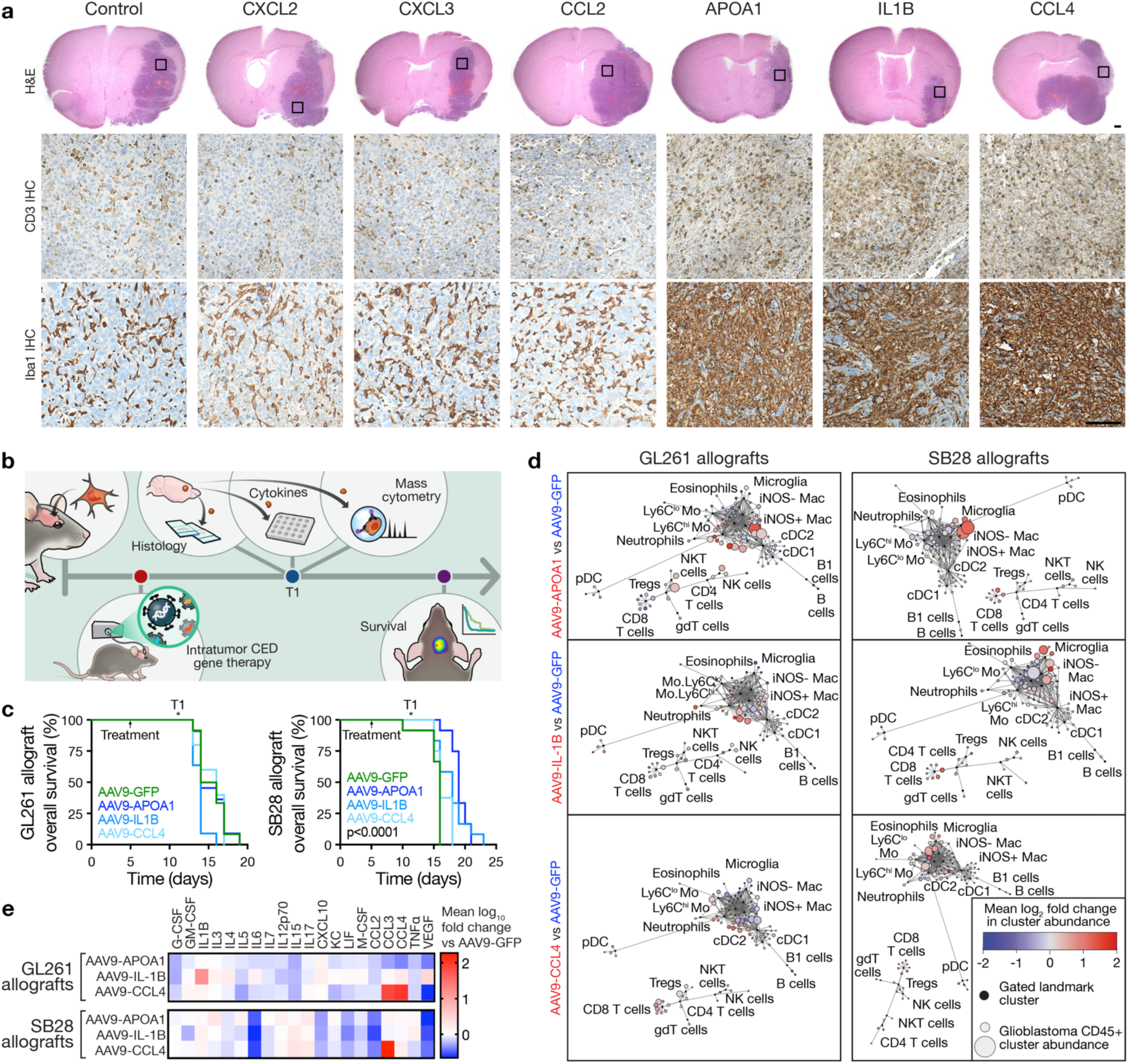
Intratumor cytokine gene therapy reprograms the glioblastoma immune microenvironment. **a**, H&E (top), CD3 IHC (middle), and Iba1 IHC (bottom) sections from intracranial glioblastoma GL261 allografts in C57BL/6J mice treated with intratumor CED of recombinant peptides associated with T cell infiltration in benign intracranial tumors (3.5-10ug recombinant peptide/mouse). Boxes show locations of IHC images. Scale bars, 1mm and 100µm. **b**, Experimental workflow for intracranial glioblastoma allograft implantation, intratumor CED gene therapy, sample collection for cellular and molecular analyses at timepoint T1, and longitudinal survival analysis. **c**, Kaplan-Meier curves for overall survival from GL261 (left, n=10 mice/condition) or SB28 (right, n=10 mice/condition) intracranial glioblastoma allografts after intratumor CED AAV9 treatments showing improved overall survival from SB28 glioblastomas with cytokine gene therapies compared to AAV9-GFP control (2x10^11^ viral genomes per mouse). Log rank tests. **d**, CD45+ immune cell CyTOF scaffold plots (n=2,291,320 cells for GL261, n=1,485,064 cells for SB28) showing unsupervised cell clusters from GL261 (left) or SB28 (right) intracranial glioblastoma allografts after intratumor CED treatments with AAV9 gene therapies vs AAV9-GFP (n=4 mice/condition), showing T cell and microglia infiltration in SB28 glioblastomas after treatment. Manually gated landmark immune cell populations (black) are annotated. **e**, Heatmap of cytokine changes from multiplexed bead assays of GL261 (top) or SB28 (bottom) intracranial glioblastoma allograft tumor lysates after intratumor CED treatments with AAV9 gene therapies (n=3 mice/condition) vs AAV9-GFP (n=3 mice) showing suppression of IL6 with all gene therapies in SB28 but not GL261 glioblastomas.

### IL-6 from tumor and non-tumor cells in the glioblastoma microenvironment increases tumor-initiating capacity and reduces survival

Re-analysis of 32,877 single-cell transcriptomes from 11 human glioblastomas^41^ (IDH-wildtype, CNS WHO grade 4) revealed that *IL-6* and *IL-6R* were expressed by tumor and diverse tumor microenvironment cells, including radial glial cells and other stem, precursor, and progenitor cells (Fig. 6a, b). These data corroborate the enrichment and correlation of stemness with IL-6-associated genes in ICI non-responders that were analyzed using spatial transcriptomics (Fig. 2). Indeed, in patient-matched human glioblastomas pre- and post-ICI (Fig. 2), spatial transcriptomic profiling demonstrated *IL-6* suppression in pre-ICI samples from ICI responders compared to pre-ICI samples from non-responders (Fig. 6c), suggesting that reduced *IL-6* may facilitate glioblastoma responses to ICI. RNA-sequencing data from the Ivy Glioblastoma Atlas Project^42^, where cells from distinct histological regions were isolated by microdissection, showed that *IL-6* expression was highest in viable pseudopalisading tumor cells adjacent to intratumor necrosis^43^ (Fig. 6d), and analysis of The Cancer Genome Atlas (TCGA) outcomes data showed that elevated *IL-6* expression was associated with reduced overall survival in glioblastoma patients (Fig. 6e).

**Fig. 6.**
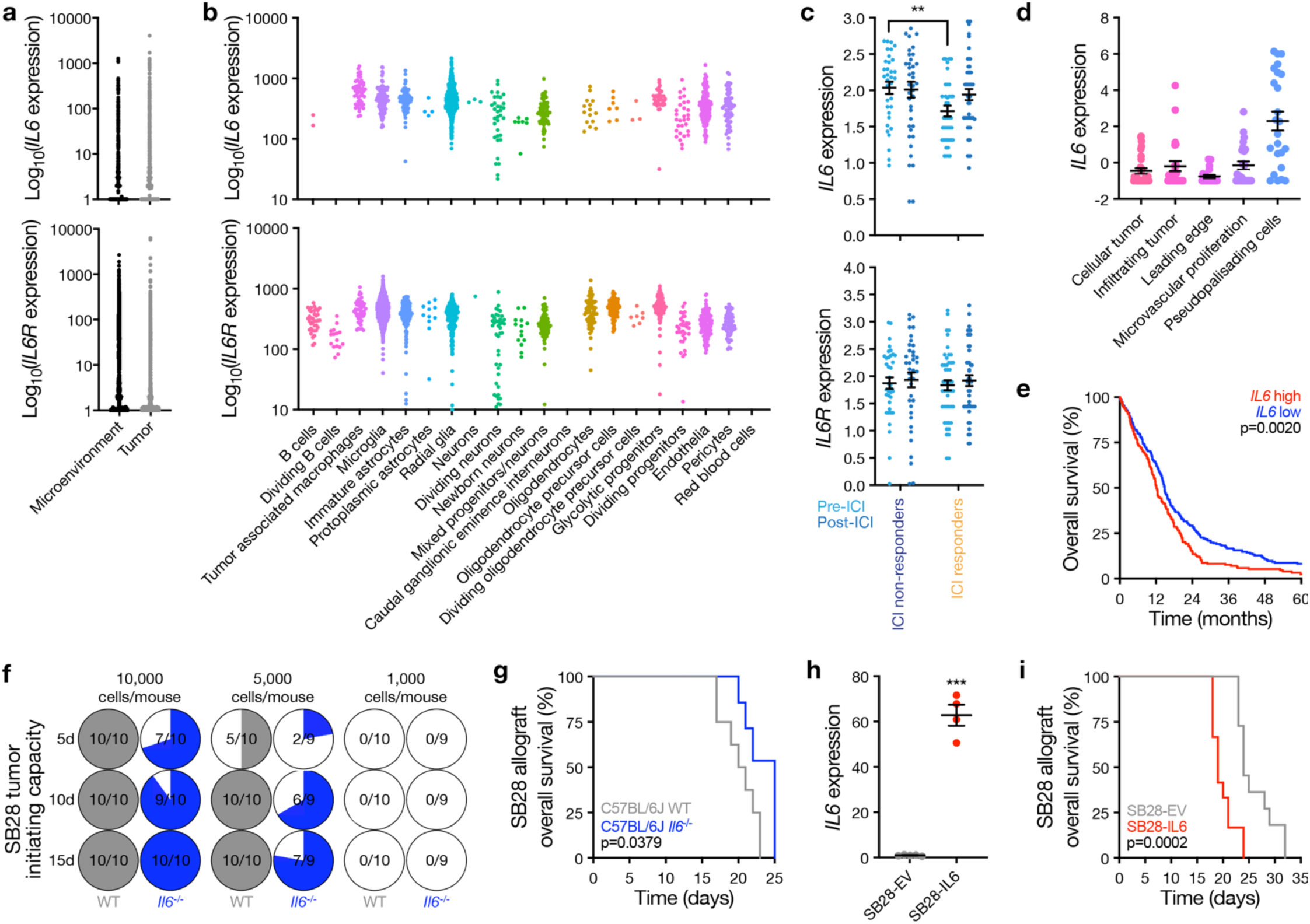
IL6 from tumor and non-tumor cells in the glioblastoma microenvironment increases tumor initiating capacity and reduces overall survival. **a** and **b**, Single-cell RNA sequencing of 32,877 cells from 11 human glioblastomas showing *IL6* and *IL6R* are expressed by tumor cells and diverse tumor microenvironment cell types in human glioblastomas. **c**, Spatial expression of *IL6* or *IL6R* from 7 patients with 14 match-paired glioblastoma samples that were resected before or after treatment with ICI showing *IL6* is suppressed in ICI responders vs ICI non-responders pre-ICI. Student’s t test, **p≤0.01. **d**, *IL6* expression across intratumor histological locations from human glioblastomas in the Ivy Glioblastoma Atlas Project (RNA sequencing from n=122 glioblastoma samples, 10 patients) showing *IL6* is enriched in viable pseudopalisading cells adjacent to regions of intratumor necrosis. ANOVA, p<0.0001. **e**, Kaplan-Meier curves for overall survival from *IL6* high (n=263 patients) vs *IL6* low (n=262 patients) human glioblastomas in The Cancer Genome Atlas. *IL6* expression was dichotomized at the mean. Log-rank test. **f**, *In vivo* intracranial tumor initiating capacity of SB28 glioblastoma allograft cells in C57BL/6J wildtype (WT) vs C57BL/6J *Il6*^-/-^ mice showing *IL6* from the microenvironment increases glioblastoma tumorigenesis. Denominators indicate number of mice at each time point. Numerators indicate number of mice with tumors as measured using intracranial bioluminescence at each time point. **g**, Kaplan-Meier curves for overall survival from SB28 intracranial glioblastoma allografts in C57J/B6 WT (n=8 mice) vs C57J/B6 *Il6^-/-^* mice (n=8 mice) showing *Il6* from the glioblastoma microenvironment reduces overall survival. Log rank test. **h**, QPCR assessment of *IL6* in SB28 cells with doxycycline-induced *IL6* overexpression compared to SB28 empty vector (EV) cells treated with doxycycline. Student’s t test, ***p≤0.0001. **i**, Kaplan-Meier curves for overall survival from SB28 intracranial glioblastoma allografts with doxycycline-induced *IL6* (n=10 mice) or EV (n=10 mice) overexpression showing *IL6* from glioblastoma tumor cells reduces overall survival. Log rank test.

Given the association between IL-6 expression and glioblastoma outcome, we tested whether IL-6 functionally contributes to glioblastoma growth in murine glioblastoma. Intracranial SB28 tumor-initiating capacity was reduced in C57BL/6J *Il-6* knockout mice compared to wild-type mice (Fig. 6f). Furthermore, survival was longer in the *Il-6* knockout mice relative to wild-type mice (Fig. 6g), which was not the case when SB28 glioblastoma was implanted into C57BL/6J *Ccl2* or *Cxcl10* knockout mice (Extended Data Fig. 6). These two cytokines that were also found to be suppressed after intratumor cytokine gene therapy treatments (Fig. 5e). Conversely, mice with intracranial SB28 that overexpressed IL-6 upon doxycycline induction were found to have significantly shorter survival than mice implanted with SB28 wild-type (Fig. 6h, i). These data reinforce the associations between IL-6 with glioblastoma stemness and reveal a functional role for IL-6 in driving glioblastoma growth.

### Combined IL-6 and PD-1 blockade reprograms the glioblastoma immune microenvironment and improves survival in preclinical models

IL-6 inhibition alleviates immunosuppression from innate immune cells in cancer^13,44,45^, and PD-1 inhibition blocks the suppression of adaptive immunity. Thus, we hypothesized that the combination of IL-6 and PD-1 blockade could improve survival from glioblastoma. To test this, we administered IL-6 and/or PD-1 blocking antibodies systemically to C57BL/6J mice with intracranial SB28 glioblastoma. Mice were monitored for survival to determine treatment efficacy, and sequential tissue samples were collected for CyTOF analysis to capture dynamic innate and adaptive cell population changes in the TIME (Fig. 7a). Combined IL-6 and PD-1 blockade modestly improved survival in mice bearing intracranial SB28, but neither antibody exerted therapeutic activity as a monotherapy (Fig. 7b). Intratumor delivery of IL-6 blockade was unable to reprogram the glioblastoma TIME or prolonging survival, even when combined with PD-1 blockade (Extended Data Fig. 7), suggesting that blockade of IL-6 from sources outside the TIME is required for treatment efficacy. CyTOF data were analyzed using Scaffold^46^ for all immune cells (Fig. 7c), and for specific T cell and myeloid cell populations using UMAP and Phenograph clustering^47^ (Fig. 7d-f). For lymphoid cells, systemic delivery of combined IL-6 and PD-1 blockade induced naïve and effector CD8+ T cell expansion in the TIME at timepoint #1 (T1, 4 days post-treatment initiation), while suppressing immunosuppressive Treg infiltration that was prominent with PD-1 blockade monotherapy (Fig. 7c-f). However, the effects from combined IL-6 and PD-1 blockade were transient, and TIME changes in CD8+ T cells did not persist at timepoint #2 (T2, 10 days post-treatment initiation). For myeloid cells, M1-like iNOS+ macrophages and monocytes increased in the TIME at T1 (Fig. 7c, f), but at T2, these changes again did not persist (Fig. 7e, f). To further interrogate interactions between adaptive and innate immune cells during combination treatment, we performed co-correlation analyses of cell clusters from SB28 treated with combined IL-6 and PD-1 blockade at T1 and T2. At T1, there were positive innate-adaptive immune correlations between CD8+ T cells, CD4+ T cells, NK cells, and activated MHCII+ monocytes, but each of these correlations diminished by T2 (Fig. 7g). Instead, CD4+ T cells and NK cells co-correlated with CD206+ macrophages at T2, suggesting co-expansion of immunosuppressive myeloid cells alongside anti-tumoral lymphocytes and innate immune cells (Fig. 7g). These data show that systemic IL-6 and PD-1 blockade modestly improves survival in mice bearing intracranial SB28 through transient reprogramming of the TIME in a model that closely resembles the human glioblastoma TIME (Fig. 4).

**Fig. 7.**
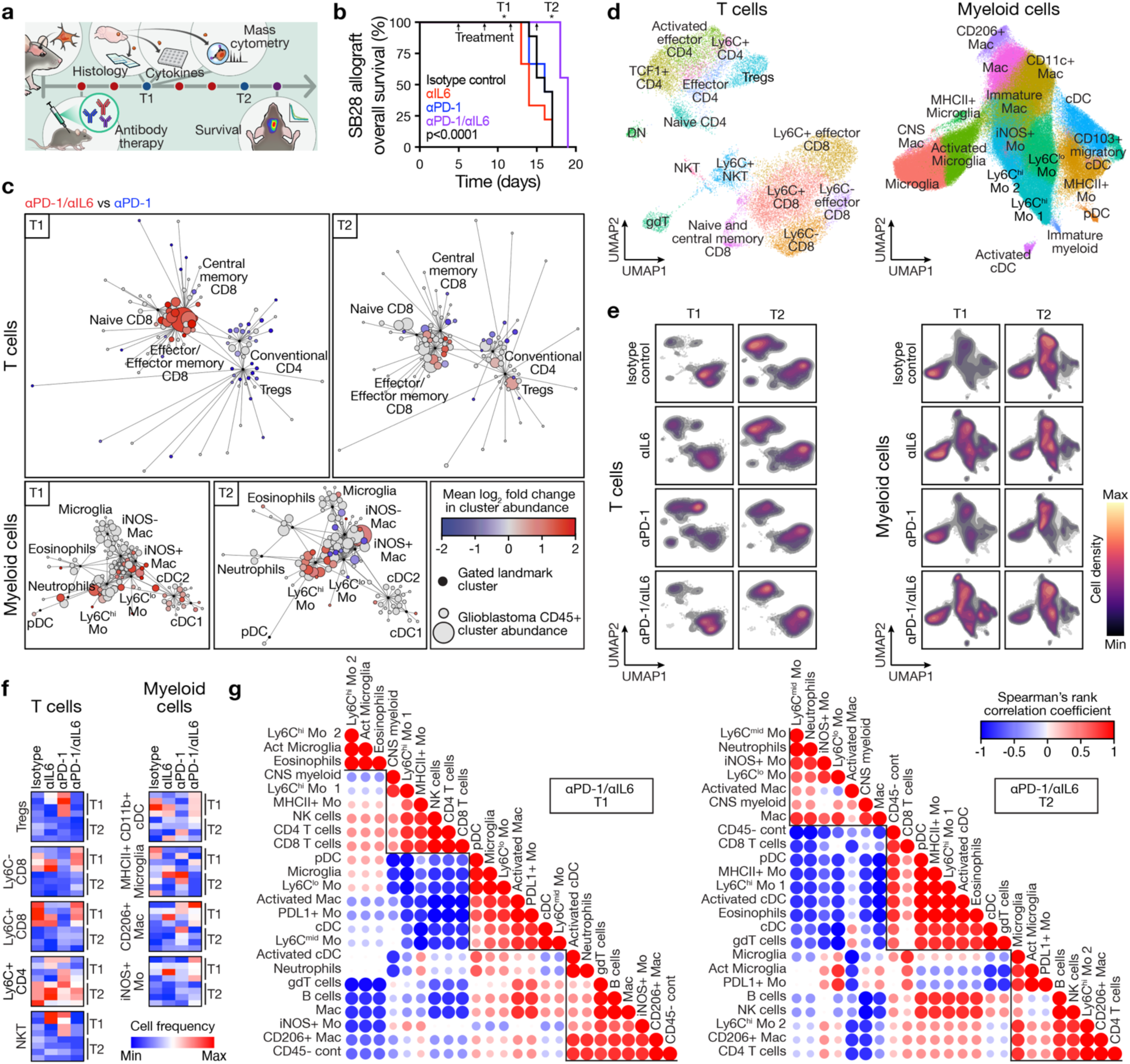
Combined IL6 blockade and immune checkpoint inhibition reprograms the glioblastoma immune microenvironment and improves overall survival. **a**, Experimental workflow for intracranial glioblastoma allograft implantation, serial systemic antibody therapy, sample collection for cellular and molecular analyses at timepoints T1 (4 days after treatment initiation) and T2 (10 days after treatment initiation), and longitudinal survival analysis. **b**, Kaplan-Meier curves for overall survival from SB28 intracranial glioblastoma allografts showing improved overall survival after systemic treatment with combined αPD-1/αIL6 therapy (250 μg biweekly i.p.) compared to monotherapy or isotype control treatments (n=8-9 mice/condition). Arrows indicate serial antibody treatments. Log rank test. **c**, CD45+ immune cell CyTOF scaffold plots of T cells (n=24,211 cells, top) or myeloid cells (n=504,571 cells, bottom) from SB28 intracranial glioblastoma allografts at T1 or T2 timepoints during combined αPD-1/αIL6 therapy vs αPD-1 monotherapy. Immune cell types in proximity to manually gated landmark immune cell populations (black) are colored when the proportion of cells increased during combined αPD-1/αIL6 therapy (red) vs αPD-1 monotherapy (blue), with q-value <0.1 by significance analysis of microarrays (SAM). At T1 there was an increase in CD8+ T cells following combination therapy while T2 was marked by an increase in monocytes and macrophages without a corresponding increase in CD8+ T cells. **d**, UMAP and Phenograph analysis of T cells (n=19,856 cells, left) or myeloid cells (n=170,000 cells, right) in the immune microenvironment in SB28 intracranial glioblastoma allografts at T1 and T2 timepoints during serial systemic antibody therapy. **e**, UMAP cell density plots for T cells (left) or myeloid cells (right) at T1 or T2 timepoints during serial systemic antibody therapy of SB28 intracranial allografts. **f**, Heatmaps of T cell or myeloid cell subset frequencies from **d** for individual mice. All cell subsets shown have Student’s t test p≤0.05 in at least one T1 vs T2 comparison across treatment conditions. Combined systemic αPD-1/αIL6 therapy increased CD8+ T cells and CD11b+ cDCs at T1. CD4+ T cells increased at T2. **g**, Spearman correlation matrices for all CD45+ immune cell subset frequencies determined from CyTOF UMAP data from **d** at T1 (left) or T2 (right) timepoints during serial systemic antibody therapy of SB28 intracranial allografts. Cluster order is determined using hierarchical clustering to identify enriched hubs (triangles) of co-correlating immune cell types. At T1, there were positive correlations between CD8+ T cells and CD4+ T cells and NK cells. At T2, there were negative correlations between CD8+ T cells and neutrophils, Ly6C^lo^ monocytes, and Ly6C^mid^ monocytes.

### Combined IL-6 and PD-1 blockade synergizes with radiotherapy to reprogram the glioblastoma TIME and significantly improve survival in preclinical models

Daily low-dose radiotherapy is the backbone of adjuvant glioblastoma treatment, but T-cells in the treatment field are particularly sensitive to the cytotoxic effects of daily ionizing radiation^48,49^. In contrast, moderate-to-high dose radiotherapy delivered over a few fractions during glioblastoma recurrence^50,51^ can have favorable immunomodulatory effects on the TIME, including MHC upregulation and lymphocyte activation^52,53^. Thus, we investigated the effects of combined IL-6 and PD-1 blockade with immunomodulatory radiotherapy (18Gy in 1 fraction). Radiotherapy was a more effective monotherapy in GL261 relative to SB28, suggesting that the relatively T cell-rich TIME in GL261 is beneficial for tumor control after immunomodulatory irradiation (Fig. 4g, 8a, Extended Data Fig. 10). CyTOF Scaffold analysis of glioblastoma showed enrichment in myeloid cells compared to the T cells and NK cells after radiotherapy (Fig. 8b). Using UMAP and Phenograph clustering of CyTOF data, radiotherapy recruited more putative immunosuppressive myeloid cells such as Ly6C^hi^ monocytes, MHCII-macrophages, and microglia to the TIME of SB28 than GL261 (Fig. 8c-d). Moreover, there was depletion of numerous T cell subsets in SB28 after radiotherapy, including naive and activated CD8+ T cells, CD4+ T cells, and ψ8T cells (Fig. 8e). Given the relative paucity of cytotoxic immune cells and immunosuppressive myeloid predominance despite radiotherapy in SB28, we hypothesized that combined IL-6 and PD-1 blockade, which increases T cell infiltration and reprograms the myeloid compartment in this preclinical model (Fig. 7), would promote immune activation in favor of tumor control. In support of this hypothesis, treatment with radiotherapy followed by combined IL-6 and PD-1 blockade prolonged survival for mice bearing intracranial SB28 compared to radiotherapy alone (Fig. 8g, Extended Data Fig. 13).

**Fig. 8.**
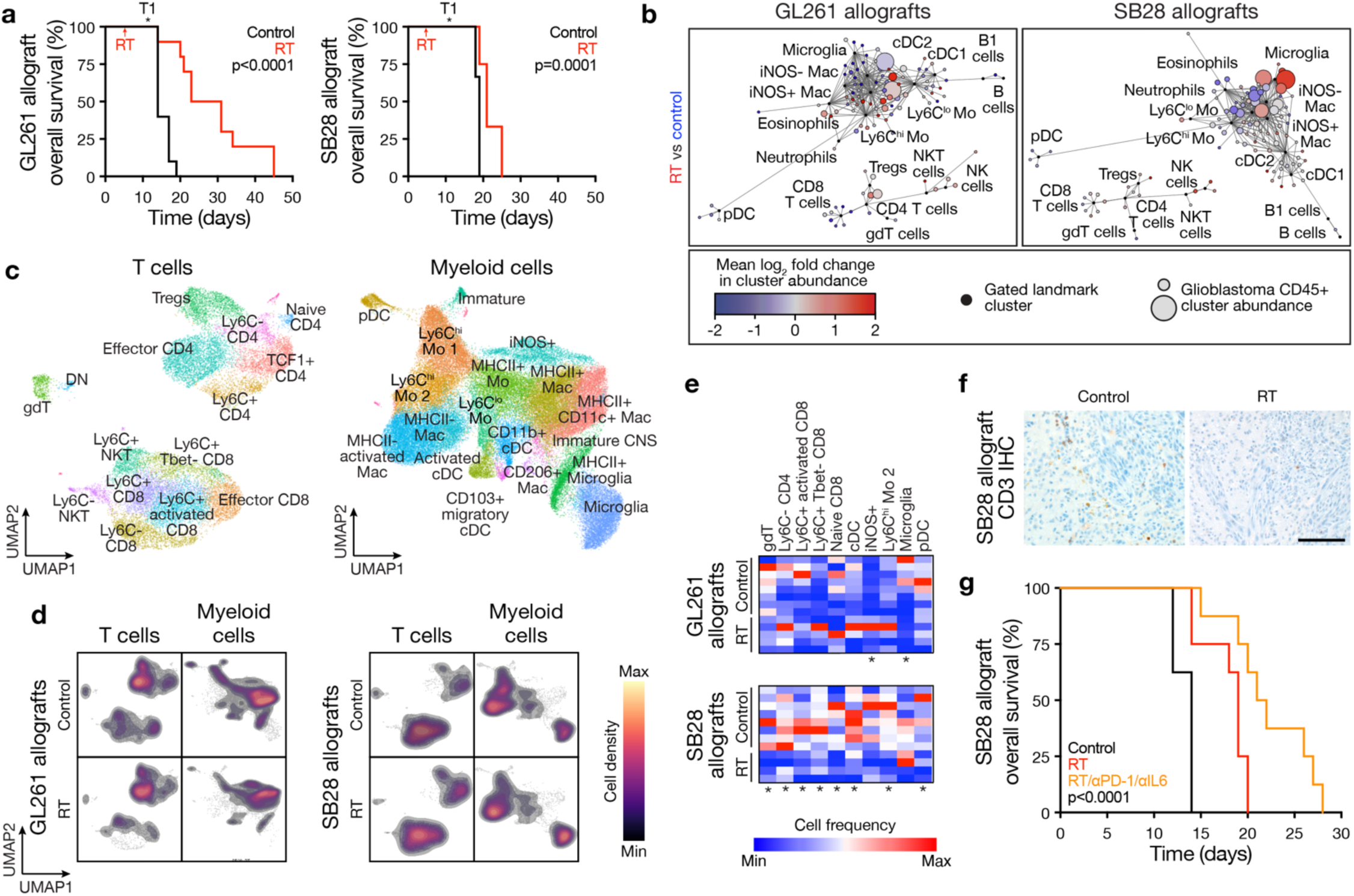
Combined IL6 blockade and immune checkpoint inhibition improves overall survival after ablative radiotherapy for glioblastoma. **a**, Kaplan-Meier curves for overall survival from GL261 (left, n=10 mice/condition) or SB28 (right, n=10 mice/condition) intracranial glioblastoma allografts after radiotherapy (RT, 18Gy/1Fx) vs control showing long-term survival from GL261 glioblastomas and short-term survival from SB28 glioblastomas after ablative RT. T1 shows sample collection timepoint for cellular and molecular analyses. Log rank tests. **b**, CD45+ immune cell CyTOF scaffold plots from GL261 (n=545,566 cells, left) or SB28 (n=506,245 cells, right) intracranial glioblastoma allografts after RT (n=4-5 mice/condition) vs control (n=4 mice) showing microglia infiltration in SB28 glioblastomas after ablative RT. Manually gated landmark immune cell populations (black) are annotated. **c**, UMAP and Phenograph analysis of T cell types (n=19,600 cells, left) or myeloid cells (n=62,500 cells, right) in the immune microenvironment of GL261 and SB28 intracranial glioblastoma allografts after RT and control (n=8 mice for control, n=9 mice for RT). **d**, UMAP cell density plots for manually gated T cells (left) or myeloid cells (right) of GL261 and SB28 intracranial glioblastoma allografts after RT vs control. **e**, Heatmap of T cell or myeloid cell subset frequencies from **c,** showing RT depletes T cells in SB28 glioblastomas. The majority of significantly changed immune cell types were identified in SB28 intracranial glioblastoma allografts. Student’s t test, *p≤0.05. **f**, CD3 IHC sections from SB28 intracranial glioblastoma allografts after RT vs control. Representative of n=3 mice/condition. Scale bar, 100µm. **g**, Kaplan-Meier curves for overall survival from SB28 intracranial glioblastoma allografts showing improved overall survival after RT vs control and after combined αPD-1/αIL6 therapy plus RT vs RT or control (n=8-10 mice/condition). Log rank test.

## Discussion

T-cell activation with ICI has been successful for multiple solid and hematological malignancies^54^, but these outcomes in glioblastoma have been discouraging and there has been no survival benefit of PD-1 blockade in large clinical trials^55^. These disappointing clinical results are likely a product of the highly immunosuppressive glioblastoma TIME, which has a paucity of effector T-cells and a predominance of myeloid cells^56^. Although IL-6 exerts pleiotropic effects on the immune system^13,20^, it has emerged as a critical cytokine for the activation of immunosuppressive myeloid cells in cancer and may influence clinical responses to ICI^57^.

Using spatial protein and transcriptomic profiling in conjunction with multiplexed sequential immunofluorescence, we first show that ICI-resistant human glioblastomas are enriched in cancer stem cell markers and immunosuppressive macrophages and myeloid cells. Our data further reveals that ICI-resistant human glioblastomas are enriched in genes that induce IL-6, such as *LGALS3,* and downstream targets of the IL-6 signaling cascade such as p-STAT3, p-AKT, p-ERK1/2, and *ANXA1*^13,28,29^. We also show that the SB28 mouse glioblastoma model closely recapitulates the TIME of human glioblastomas and that multiple immune-activating gene therapies decrease intratumoral IL-6 levels in SB28 tumors. In human glioblastoma, IL-6 levels are endogenously lower in patients who achieve histological or radiographic responses to ICI, and ICI-resistant glioblastoma has elevated stem cell markers (SOX2, SOX4, SOX13, PTPRZ1), which can be driven by IL-6^21,58^. Blocking IL-6 has been previously shown to suppress the growth of subcutaneously implanted human glioma stem cell-derived xenografts in immunocompromised mice^18^. We now show that systemic blockade of IL-6 and anti-PD-1 prolongs survival in intracranial SB28 immunocompetent mice with the corresponding expansion of CD8+ T-cells and suppression of Treg infiltration. Our data also showed that these cellular changes in the TIME are transient, leading to treatment failure^59^. We find that ablative radiotherapy, which increases microglia and decreases T-cells in the TIME, is more effective when combined with concurrent IL-6 and ICI blockade. Together, these results suggest that IL-6 blockade can be combined with ICI and radiotherapy to enhance treatment responses in human glioblastoma patients. In support of this hypothesis, IL-6 receptor-neutralizing antibodies are being tested with PD1 blockade and re-irradiation in a Phase 2 clinical trial for recurrent glioblastoma (NCT04729959).

Although these results are encouraging, the changes in the glioblastoma TIME are transient after combined IL-6 and ICI blockade, suggesting that the immunosuppressive TIME is durable and persistent despite treatment and likely necessitating a sustained treatment schedule. Indeed, our data and data from other investigators suggest that the survival benefit from IL-6 plus ICI blockade is modest, but when combined with additional agents, such as CD40 stimulation or radiotherapy, this therapeutic strategy can durably prolong survival in glioblastoma preclinical models^15^. The interaction between IL-6 in the TIME and cancer stem cell enrichment warrants further investigation as a potential therapeutic target in glioblastoma, as the relationship between IL-6 signaling and cancer stem cell growth has been established in other solid malignancies^60^ and is likely related to STAT3^61^.

## Methods

### Inclusion and ethics

This study complied with all relevant ethical regulations and was approved by the University of California San Francisco (UCSF) Institutional Review Board (IRB #15-17500). As part of routine clinical practice, all patients in this study signed a written waiver of informed consent to contribute de-identified tissue for research.

### Human glioblastoma patient clinical data, histology, and microscopy

The study cohort of human patients was comprised of 14 matched-paired IDH-wildtype glioblastoma samples, CNS WHO grade 4, from 7 patients who were treated with immune checkpoint inhibition (ICI) after initial resection and standard-of-care chemoradiotherapy and eventually went on to re-resection of recurrent glioblastoma at UCSF. Patients were classified as ICI responders versus ICI non-responders based on previously defined criteria^22,23^: (1) tissue sampled during surgery after ICI immunotherapy showing robust reactive immune infiltrate with scant or no tumor cells, or (2) tumor size stability or decreasing tumor size over at least 6 months from initiation of ICI immunotherapy on magnetic resonance imaging (MRI). Tissue from a separate cohort of 8 human IDH-wildtype CNS WHO grade 4 glioblastoma patients who underwent resection in the absence of ICI therapy were collected for mass cytometry by time-of-flight (CyTOF) cell profiling and analysis. Patient demographics, treatments, and clinical outcomes were recorded from the electronic medical record (Supplementary Table 1). MRI studies were reviewed to define glioblastoma locations and clinical outcomes. Pathologic examination of the entire cohort was performed by a board-certified neuropathologist (C-H.G.L) to assess for histological or molecular heterogeneity and to select regions of interest (ROI) with the highest cell viability for spatial protein profiling and spatial transcriptomic analyses.

### Targeted next-generation DNA sequencing and analysis

Targeted DNA sequencing was performed on all human samples using the UCSF500 next-generation DNA sequencing panel, as previously described^62^. In brief, this capture-based assay targets all coding exons of 479 cancer-related genes, select introns, and upstream regulatory regions of 47 genes to enable the detection of structural variants, such as gene fusions, and DNA segments at regular intervals along each chromosome to enable genome-wide copy number and zygosity analyses, with a total sequencing footprint of 2.8 Mb (Supplementary Table 2). Multiplex library preparation was performed using the KAPA Hyper Prep Kit (Roche, cat# 07962355001). Hybrid capture of pooled libraries was performed using a custom oligonucleotide library (Nimblegen SeqCap EZ Choice). Captured libraries were sequenced as paired-end reads on an Illumina NovaSeq 6000 at >200x coverage for each sample. Sequence reads were mapped to the reference human genome build GRCh37 (hg19) using the Burrows-Wheeler aligner (v0.7.17). Recalibration and deduplication of reads were performed using the Genome Analysis Toolkit (v4.3.0.0). Coverage and sequencing statistics were determined using Picard (v2.27.5), CalculateHsMetrics, and CollectInsertSizeMetrics. Single nucleotide variant and small insertion/deletion mutation calling were performed with FreeBayes, Unified Genotyper, and Pindel. Large insertion/deletion and structural alteration calling were performed with Delly. Variant annotation was performed with Annovar. Single nucleotide variants, insertions/deletions, and structural variants were visualized and verified using Integrative Genome Viewer (v.2.16.0). Genome-wide copy number and zygosity analyses were performed using CNVkit and visualized using NxClinical (Biodiscovery, v6.0).

### Spatial protein profiling and analysis

Spatial protein profiling was performed on formalin-fixed paraffin-embedded (FFPE) tumor sections using the NanoString GeoMx Digital Spatial Profiler at the UCSF Laboratory for Cell Analysis Genome Core. Glioblastoma FFPE sections cut at 5 um thickness on charged slides were labeled with morphology markers FITC (Invitrogen, cat# S7575) and CD3 (Origen, cat# UM500048, clone UMAB54), as well as a multiplexed cocktail of 78 oligo-conjugated antibodies (Supplementary Table 3) using human protein panel modules generated at NanoString Technologies. Incubation with morphology marker CD3 (1:40) and oligo-conjugated antibodies was performed at 4°C in a humidity chamber over 12 hours. Subsequent incubation with morphology marker FITC (1:10) was performed at room temperature over 20 minutes. H&E stained whole slide images were overlayed on fluorescent projections and 200μm ROI were annotated using the GeoMx Digital Spatial Profiler (Nanostring) based on viable tumor tissue regions to avoid confounding results from regions of intratumor necrosis by a board-certified neuropathologist (C-H.G.L). ROI for spatial protein profiling overlapped with those selected for spatial transcriptomic sequencing, as described below. Photocleavable oligonucleotides were released from ROI using ultraviolet cleavage, aspirated tags were hybridized to optical barcodes and processed using the NanoString nCounter Analysis System. Barcodes were normalized with internal spike-in controls and then normalized against housekeeping genes.

### Spatial transcriptomic sequencing and analysis

Spatial transcriptomic profiling was performed on FFPE sections using the Nanostring GeoMx Digital Spatial Profiler at the UCSF Laboratory for Cell Analysis Genome Core. Glioblastoma FFPE sections cut at 5 um thickness on charged slides were labeled with morphology markers FITC and CD3 as described above and hybridized with an RNA probe mixture targeting approximately 18,000 protein-coding genes using the manufacture protocol. Hybridization with the RNA probe mixture was performed at 37°C for 16 hours. Subsequent incubation with morphology markers FITC (1:10) and CD3 (1:40) was performed at room temperature for 1 hour. H&E stained whole slide images were overlayed on fluorescent projections and 200μm ROI were annotated using the GeoMx Digital Spatial Profiler (Nanostring) based on viable tumor tissue regions to avoid confounding results from regions from intratumor necrosis by a board-certified neuropathologist (C-H.G.L). Some ROI for spatial transcriptomic sequencing overlapped with those selected for spatial protein profiling, as described above, and additional ROIs for spatial transcriptomic sequencing were chosen based on viable tissue by C-H.G.L. Photocleavable oligonucleotides were released from regions of interest using ultraviolet light, aspirated tags were hybridized to optical barcodes, sequencing libraries underwent polymerase chain reaction per manufacture recommended protocol, and pooled captured libraries were sequenced on an Illumina NovaSeq 6000. Quality control on resulting ROI was performed according to manufacturer recommendations, which resulted in 173 out of 190 ROIs passing quality control. Data were normalized to the 3^rd^ quartile (Q3 normalization) per manufacturer recommendations. COMBAT batch correction was used to correct for slide batch effects^63^. Q3 normalized, COMBAT batch corrected, and log2 transformed counts were used in subsequent analyses. Differential gene expression was performed using methods recommended by the manufacturer^64^, via a linear mixed-effect model in R (v3.6.2) and the lmer function in the lmerTest package (v3.1-3) with default settings to generate a random slope (by patient) and random intercept (by slide) linear mixed-effect model. P-values reported for differentially expressed genes were corrected for false discovery using the Benjamini-Hochberg method. Heatmaps were generated using the pheatmap function of the pheatmap package (v1.0.12), with default settings (clustering_method = “average”, clustering_distance_rows = “correlation”).

### Pathway enrichment and visualization analysis

Gene Set Enrichment Analysis (GSEA, v.4.3.2) was performed to identify pathways enriched in differentially expressed genes. The gene rank scores were calculated using the formula: SIGN(log2FC) × −log10(p-value). Pathways were delineated using Human_GOBP_AllPathways_no_GO_iea_July_03_2023_symbol.gmt, which is periodically updated and maintained by the Bader laboratory. Positive and negative enrichment profiles were achieved through 2000 permutations. The gene set size was constrained between 10 to 500 for the analysis. Pathway analysis results were visualized with the EnrichmentMap App (v.3.3.6) using Cytoscape (v.3.10.0). Parameters set for nodes included an FDR q-value of less than 0.05, a p-value of less than 0.05, and nodes that shared gene overlaps with a Jaccard + Overlap Combined (JOC) threshold of 0.375. Such nodes were interconnected with a blue line (edge) to formulate network maps. Clusters of analogous pathways were pinpointed and labeled utilizing the AutoAnnotate app (v.1.4.1) in Cytoscape. This app incorporates a Markov Cluster algorithm, connecting pathways by mutual keywords in their description. The resulting groups of pathways were designated as the major pathways in circles.

### Multiplexed sequential immunofluorescence (seq-IF) and microscopy

The multiplex panel included the following markers (antibody information can be found in Supplementary Table 5): CD31, GFAP, CD4, CD8, CD11c, CD68, CD163, P2RY12, p-STAT3, SOX2, PTPRZ1, p-ERK1/2, PD-1, CTLA-4, LAG-3, TIM-3, IL-6R. The antibodies were validated using either immunohistochemistry and/or immunofluorescence staining. The optimal concentration and the best signal/noise ratio was identified by sequential dilution. The secondary Alexa fluorophore 555 (ThermoFisher Scientific) and Alexa fluorophore 647 (ThermoFisher Scientific) were used at 1:200 and 1:400 dilution, respectively. The optimizations and full runs have been previously ^65,66^. The elution step lasted 2 min for each cycle at 37°C and quenching lasted for 30 sec. Imaging exposure time was set at 4 min for all primary antibodies, except for IL-6R at 8 mins, and secondary antibodies at 2 min. The image registration was performed after the staining using COMET Control Software. Markers were pseudocolored for visualization of markers in the Viewer from Lunaphore and quantified using Visiopharm software.

### Cell culture and molecular biology

Luciferase-expressing mouse glioblastoma GL261 and SB28 cell lines (passage number 20-28) were cultured in DMEM/F12 (ThermoFisher Scientific, cat# 11320033) containing 4.5 g/l glucose and 10% FBS. Cells were maintained in a humidified incubator at 5% CO2 at 37°C and passaged when 70-80% confluent (every 2-4 days). The cell lines were routinely confirmed to be negative for mycoplasma infection. Patient-derived human glioblastoma xenograft cells (GBM6) were maintained *in vivo* and were provided by the UCSF Brain Tumor Preclinical Core.

To generate a doxycycline-inducible IL-6 lentiviral vector, gene fragment blocks representing the coding sequencing of *Il6* (NM_001314054.1) were purchased from IDT DNA Technologies. The coding sequence was appended with a 3x-FLAG tag, a Kozak sequence, and restriction enzyme sites for EcoRI and BamHI with a minimum of at least 6 nucleotides preceding and proceeding the cut sites. The dox-inducible lentiviral vector pLV.TreGS and the gene fragment blocks were digested for 2 hours at 37°C with EcoRI (New England Biolabs, cat# R0101) and BamHI (New England Biolabs, car# R0136), run on a 1-2% agarose gel at 120V, excised using a razor blade, and gel extracted (Qiagen, cat# 28704). The molar ratios of the vector and insert were varied using the https://nebiocalculator.neb.com/#!/ligation calculator. Ligation was performed overnight at 16°C and plasmids were transformed into Stbl3 competent cells. The completed construct was sequenced to ensure proper ligation prior to experimentation.

Lentivirus packaging of the doxycycline-inducible IL-6 construct was performed by transient transfection in human embryonic kidney (HEK) 293T cells maintained in antibiotic free Advanced DMEM (Gibco, cat# 12491) with 3% FBS and 1% CTS GlutaMAX (Gibco, cat #A1286001). The transfection mixture of DNAs was comprised of 6 µg construct, 2 µg pMD2.G (Addgene, cat# 12259), and 6 µg psPAX2 (Addgene, cat# 12260), and was diluted into 850 µL of Opti-MEM (Gibco, cat# 31985070) with 40µl of TransIT-Lenti transfection reagent (Mirus, cat# MIR6606), followed by brief vortexting and incubation for 20 minutes at room temperature. HEK293T cells in a 10cm dish were refreshed with 9ml of culture medium, and the transfection mixture was added dropwise onto the cells, followed by 1x Viralboost reagent (Alstem, cat# VB100). The viral particles were harvested and pooled at 48 hrs and 72 hrs post transfection. SB28 cells were transduced with viral supernatant and 8 µg/ml polybrene (EMD Millipore, cat# TR-1003) after filtration through a 0.45µm PVDF syringe filter (Genclone, cat# 25-242) to remove any residual HEK293T cells. A stably polyclonal cell line was selected using 1 µg/ml puromycin (Sigma-Aldrich, cat# P8833).

Construct expression was induced with doxycycline *in vitro* at 2 µg/mL and *in vivo* at 2mg/mL in drinking water (Sigma-Aldrich, cat# D9891). For *in vivo* induction, water was changed every 2-3 days. Expression was confirmed using quantitative reverse-transcriptase polymerase chain reaction (QPCR). To do so, RNA was extracted from SB28 cells using RNeasy Plus Mini Kit (Qiagen, cat# 74134) and cDNA was synthesized using the iScript cDNA Synthesis Kit (Bio-Rad, cat# 1708891). Target genes were amplified using PowerUp SYBR Green Master Mix and a QuantStudio 6 thermocycler (ThermoFisher Scientific). *Il6* expression (sense: 5’-GAGGATACCACTCCCAACAGACC-3’, antisense: 5’-AAGTGCATCATCGTTGTTCATACA-3’) was calculated using the DDCt method, with normalization to *GAPDH* (sense: 5’-ATGGGGAAGGTGAAGGTCG-3’, antisense: 5’-GGGGTCATTGATGGCAACAATA-3’).

### Mouse models, treatments, and measurements

C57BL/6J (IMSR_JAX:000664), *Il6^-/-^* (IMSR_JAX:002650), *Ccl2^-/-^* (IMSR_JAX:004434), and Cxcl10*^-/-^* (IMSR_JAX:006087) mice were purchased from The Jackson Laboratory. Athymic Nu/Nu mice were purchased from Inotiv. Experimental mice were female and randomly assigned to different treatment groups at 6 to 8 weeks of age. Experimental and control animals were co-housed. All animal experiments were performed in full compliance with animal protocols approved by the University of California San Francisco Institutional Animal Care and Use Committee (IACUC).

Female 6-8 week-old C57BL/6J mice were implanted with glioma cells intracranially into the right caudate/frontal lobe at a depth of 3mm approximately 3mm lateral to bregma. For all efficacy experiments using non-gene therapy treatments in SB28 models, tumor implantation was performed with 3 x 10^4^ cells unless indicated otherwise. A larger number of SB28 and GBM6 cells (3 x 10^5^) was used to ensure sizeable tumors for convection-enhanced delivery for gene therapy treatments. For efficacy experiments in GL261 models, tumor implantation was performed with 3 x 10^5^ cells. Mice were anesthetized with a mixture of inhaled isoflurane (Sigma-Aldrich, cat#26675-46-7), ketamine (Warner-Lambert), and xylazine (Sigma-Aldrich, cat#X1126) for tumor implantation, treatment, or bioluminescence monitoring. Mice were monitored daily and euthanized at veterinary-stipulated endpoints (15% weight loss and/or other clinical signs or symptoms of intracranial tumor, such as circling, inability to maintain an upright position, spasticity, seizures, paresis, or inability to ambulate). Standard post-surgery care and analgesia were provided in accordance with the IACUC-approved protocol. Mice were randomly assigned to treatment or control arms 5 days after tumor implantation.

For gene therapy treatments, 2 x 10^11^ v.g. of each gene therapy virus was delivered via convection-enhanced delivery in a total volume of 290 μl of saline over 15 minutes using the same burr hole and coordinates as tumor cell implantation. All gene therapy vectors were constructed by inserting the coding sequence for GFP or mouse cytokines into the AAV-CAG vector, which facilitates gene expression from AAV serotype 9 virus (capsid from AAV9 and ITR from AAV2) using a hybridized promoter comprised of the cytomegalovirus (CMV) early enhancer element and the chicken beta-actin promoter. Antibodies for *in vivo* experiments were: 250 μg of aPD-1 clone RMP1-14 (Bio X Cell, cat# BP0146), 250 μg of aIL-6 clone MP5-20F3 (Bio X Cell, cat# BE0046), and mouse IgG2b isotype control clone MPC-11 (Bio X Cell, cat# BE0086). Each antibody treatment was delivered in 100μL PBS using intraperitoneal injection beginning 5 days after tumor implantation. For CED of aIL-6, the maximal soluble dose was delivered in a total volume of 290 μl of saline over 15 minutes in the same manner as the gene therapy treatments described above. For radiation treatments, anesthetized mice were placed on a platform under a cesium-137 source and shielded to limit radiation exposure to a narrow 1cm horizontal beam positioned over the skull. 18Gy was delivered in a single fraction with a dose determined using previously described methods^67^.

### Histology, light microscopy, immunohistochemistry, and fluorescence microscopy

Immunohistochemical stains for T cells (CD3) and macrophages (CD68 and Iba1) were performed on whole slide sections using standard approaches. After fixation, tissue was processed, embedded in paraffin, sectioned (5 μm), and stored at -20°C prior to use. Slides were stained with hematoxylin & eosin (H&E) or immunostained using a Discovery Ultra autostainer (Ventana Medical Systems, Inc., USA). In brief, following antigen retrieval with CC1 (Ventana Medical Systems, Inc., USA) for 32 minutes, sections were stained with CD3 antibody (NB600-1441, Novus Biologicals LLC, clone SP7, rabbit monoclonal, dilution 1:100) or CD68 antibody (MO814, Agilent Dako, clone KPI, mouse monoclonal, 1:100), or Iba1 antibody (019-19741, Fujifilm Wako Chemicals U.S.A. Corporation, rabbit polyclonal, 1:500). Detection was performed using the UltraView Universal DAB kit on the Ventana Benchmark Platform (Ventana Medical Systems). Light microscopy assessment was performed on an Olympus BX43 microscope with standard objectives. Light microscopy images were obtained and analyzed using the Olympus cellSens Standard Imaging Software package (v1.16). Fluorescence microscopy was performed on a Zeiss LSM 800 confocal laser scanning microscope with Airyscan.

### Mass cytometry by time-of-flight (CyTOF) cell profiling and analysis

All mass cytometry antibodies and staining concentrations are listed in Supplementary Table 6. The Maxpar antibody conjugation kit comprised of MCP9 (Fluidigm, cat#201106A) for Cadmium or X8 (Fluidigm, cat# 201141A) for non-Cadmium metals was used to prepare primary conjugates of mass cytometry antibodies according to the manufacturer’s recommended protocol, as previously described^68^. Antibodies were stored at 4°C and each antibody clone was titrated to optimal staining concentrations using primary mouse or human samples, respectively.

Tumor-bearing brains were harvested from mice after CO2 inhalation euthanasia and were homogenized with a syringe pusher over a 70μm filter in P^69^BS/EDTA (5 mM final). All samples were processed on ice with buffers at 4°C. Tumors were finely minced and digested in RPMI-1640 with 4mg/ml collagenase IV (Worthington, cat# LS004186) and 0.1 mg/ml DNaseI (Sigma-Aldrich, cat# 69182) in a scintillation vial with a micro stir bar for 30 minutes at 37°C. After digestion, cells were filtered twice over 70μm filters with PBS/EDTA. Tissue samples were centrifuged at 500g for 5 minutes at 4°C, resuspended in PBS/EDTA, and mixed 1:1 with PBS/EDTA containing 100mM cisplatin (Enzo Life Sciences, cat# ALX-400-040) for 60 seconds before quenching 1:1 with PBS/EDTA/BSA (5 mM final). Cells were centrifuged again, resuspended in PBS/EDTA/BSA at a density between 1 x 10^6^ and 1 x 10^7^ cells/mL, fixed for 10 minutes at room temperature using 1.6% formaldehyde, washed twice with PBS/EDTA/BSA, and frozen in 100ul buffer at -80°C.

2 x 10^6^ cells from each tumor were barcoded with distinct combinations of stable Pd isotopes in 0.02% saponin in PBS as previously described^69^. Cells were washed twice with cell-staining media (PBS with 0.5% BSA and 0.02% NaN3) and combined into a single 15 mL conical tube. For staining, cells were resuspended in cell-staining media, blocked with FcX (Biolegend) for 5 minutes at room temperature, then incubated for 30 minutes at room temperature on a shaker with surface marker antibody cocktail in 1mL total volume. Cells were washed in cell-staining media then permeabilized with methanol for 10 minutes at 4°C, then washed again in cell-staining media. Cells were then incubated with intracellular antibody cocktail in the same volume as surface markers for 30 minutes at room temperature on a shaker. Cells were washed in cell-staining media then incubated with 1mL PBS containing 1:5000 Iridium intercalator (Fluidigm) and 4% final formaldehyde. Cells were maintained in this final buffer for 2 days prior to sample acquisition on a CyTOF 2 mass cytometer (Fluidigm) using Maxpar Cell Acquisition Solution. 1-5 x 10^5^ cells were acquired per each sample. Data normalization was performed as previously described using EQ Four Element Calibration Beads (Fluidigm) using Normalizer v.0.3 (Nolan Lab). All mass cytometry files were normalized together using the mass cytometry data normalization algorithm, which uses the intensity values of a sliding window of bead standards to correct for instrument fluctuations over time and between samples. After normalization and debarcoding of files, singlets were gated by event length and DNA. Live cells were identified by cisplatin negative cells.

Manually gated cell subsets were determined by gating strategy shown in Extended Data Figs. 3, 4, 8, 9, 11 and 12. Principal component analysis was performed for reach human and mouse tumor sample using cell frequencies as percentages of live leukocytes. Scaffold maps were generated as previously described using the open-source Statistical Scaffold R package available at github.com/SpitzerLab/statisticalScaffold^68^. Landmark reference nodes were gated from peripheral blood mononuclear cells, while unsupervised clusters were generated using Clustering Large Applications (CLARA) clustering from samples pooled together with each sample contributing an equal number of cells. For uniform manifold approximation and projection (UMAP) and Phenograph (v0.99.1) analyses of cell subsets, equal numbers of T cells or non-polymorphonuclear myeloid cells were extracted from each sample. For T cells, cells were gated using CD45 and CD3 expression from each sample. For non-polymorphonuclear myeloid cells, neutrophils and eosinophils were excluded to focus our analyses on cell subsets that were changing the most in the human glioblastoma samples. The myeloid cells were extracted by selecting clara clusters associated with myeloid cell landmark nodes from the Scaffold analysis. UMAP analysis was performed on ArcSinh (cofactor=5) transformed protein expression values on equal numbers of cells from each sample by randomly subsampling cells with parameters min.dist=1.0. UMAP dimensionality reduction was performed using all available protein markers except Ki-67. Phenograph was used to compute cluster assignment, and cluster identities were manually assigned using marker feature plots and hierarchical clustering of median marker intensities for each cluster. For pairwise correlation matrix analysis, cell cluster frequencies (percent of singlets) for each condition were correlated pairwise using Spearman and the R values were determined. Cell clusters were then displayed using hierarchical ordering (hclustward2 using corrplot package in R).

### Multiplexed cytokine assays

For quantification of cytokines from intracranial glioblastomas, minced tumors were lysed using RIPA buffer containing protease inhibitor (Cell Biolabs, cat# AKR-190) on ice for 10 minutes, followed by centrifugation at 14,000rpm for 10 minutes at 4°C. Supernatant containing cell lysate was collected and normalized to the same using the Bio-Rad Protein Assay (cat# 5000006). For quantification of cytokines from blood plasma, freshly isolated peripheral blood was centrifuged for 10 minutes at 4°C at 1000g, and 100μL of supernatant plasma was mixed 1:1 with 100μL of PBS/EDTA. For multiplexed cytokine quantification, tumor lysate and plasma samples were submitted for the 44-plex mouse cytokine/chemokine discovery assay array (MD44) at Eve Technologies 450.

### Single-cell RNA sequencing, Ivy Glioblastoma Atlas Project (GAP), and The Cancer Genome Atlas (TCGA) data

All single-cell RNA sequencing data (32,877 cells, n=11 human glioblastomas) were re-analyzed for *IL6* and *IL6R* expression^41^. The Ivy GAP RNA sequencing data and sample annotation containing anatomic location were downloaded from the Ivy GAP website (https://glioblastoma.alleninstitute.org/static/download.html)^70^. TCGA data were processed and visualized using the GlioVis portal (https://gliovis.bioinfo.cnio.es)^71^.

### Statistics

All experiments were performed with independent biological replicates and repeated, and statistics were derived from biological replicates. Biological replicates are indicated in each figure panel or figure legend. No statistical methods were used to predetermine sample sizes, but sample sizes in this study are similar or larger to those reported in previous publications. Data distribution was assumed to be normal, but this was not formally tested. Investigators were blinded to conditions during clinical data collection and analysis of mechanistic or functional studies. Bioinformatic analyses were performed blind to clinical features, outcomes, or molecular characteristics. The clinical samples used in this study were retrospective and nonrandomized with no intervention, and all samples were interrogated equally. Thus, controlling for covariates among clinical samples was not relevant. Cells and animals were randomized to experimental conditions. No clinical, molecular, or cellular data points were excluded from the analyses. Lines represent means, and error bars represent standard error of the means. Results were compared using Student’s t-tests. Log-rank tests, and ANOVA, which are indicated in figure panels or figure legends alongside approaches used to adjust for multiple comparisons. In general, statistical significance is shown using asterisks (*p<=0.05, **p<=0.01, ***p<=0.0001), but exact p*-*values are provided in figure panels or figure legends whenever possible.

### Reporting summary

Further information on research design is available in the Nature Research Reporting Summary linked to this article.

## Data availability

CyTOF data that support the findings of this study have been deposited to Dryad (https://doi.org/10.5061/dryad.c59zw3rkm). Publicly available TCGA and Ivy Glioblastoma Atlas Project data were used in this study. Source data are provided with this paper.

## Code availability

No custom software, tools, or packages were used. The open-source software, tools, and packages used for data analysis in this study are referenced in the methods where applicable and include AutoAnnotate (v1.3.5), Burrows-Wheeler aligner (v0.7.17), Cytoscape (v3.7.2), EnrichmentMap (v.3.3.6), DESeq2 (v1.36.0), FASTQC (v0.11.9), featureCounts (v2.0.0), Genome Analysis Toolkit (v4.3.0.0), GSEA (v4.3.2), Picard (v2.27.5), R (v3.6.1 and v4.2.1), RStudio (v2022.07.02), Seurat (v4.3.0), ImerTest (v3.1-3), pheatmap (v1.0.12), Phenograph (v0.99.1), Scaffold (v0.1), and corrplot (v0.92).

## Acknowledgements

The authors thank Anny Shai and the staff of the UCSF Brain Tumor Center Biorepository and Pathology Core (NIH P50 CA097257), Emmanuella Zacco and the staff of the UCSF Laboratory for Cell Analysis Genome Core, Eric Chow and the staff of the UCSF Center for Advanced Technology, Nicole Paulk for gene therapy vector generation, Ken Probst and Noel Sirivansanti for Illustrations, and the members of the Raleigh lab for guidance and edits. This study was supported by NIH grant T32 CA151022, the American Society of Clinical Oncology Young Investigator Award, the American Association of Neurological Surgeons Neurosurgery Research and Education Foundation, the Chan-Zuckerberg Biohub Physician-Scientist Fellowship, and the Glioblastoma Foundation Neil Peart Neurosurgery Research Fund to JSY, the Chan-Zuckerburg Biohub Physician-Scientist Fellowship and NIH grant K12 CA260225 to WCC, NIH grants R01 NS120547, and P50 CA221747 to ABH, and NIH grants R01 CA262311 and U19 CA964338 to DRR.

## Author contributions statement

All authors made substantial contributions to the conception or design of the study; the acquisition, analysis, or interpretation of data; or drafting or revising the manuscript. All authors approved the manuscript. All authors agree to be personally accountable for individual contributions and to ensure that questions related to the accuracy or integrity of any part of the work are appropriately investigated and resolved and the resolution documented in the literature. JSY, NWC, and DRR conceived and designed the study. JSY performed mouse experiments and analyzed human data with MPN and performed mass cytometry by time-of-flight cell profiling and multiplexed cytokine assays with NWC. CHGJ and KM performed spatial protein profiling and transcriptomic experiments and analyzed histological and immunohistochemical data. Mass cytometry by time-of-flight and sequencing data were analyzed NWC, WCC, NZ, and DRR. HN performed multiplexed sequential immunofluorescence under supervision by ABH. KS, VD, TCC, and AC performed molecular biology experiments. JJP designed and performed histological and immunohistochemical experiments. AB provided single-cell RNA sequencing data and analyses. TO designed and performed mouse experiments with KS. MKA, MSB, and JWT provided patient data. JLD supervised gene therapy vector development. The study was supervised by ABH, MSB, NB, MHS, and DRR. The manuscript was prepared by JSY, NWC, and DRR with input from all authors.

## Tables

Not applicable

---

**Extended Data Fig. 1.**
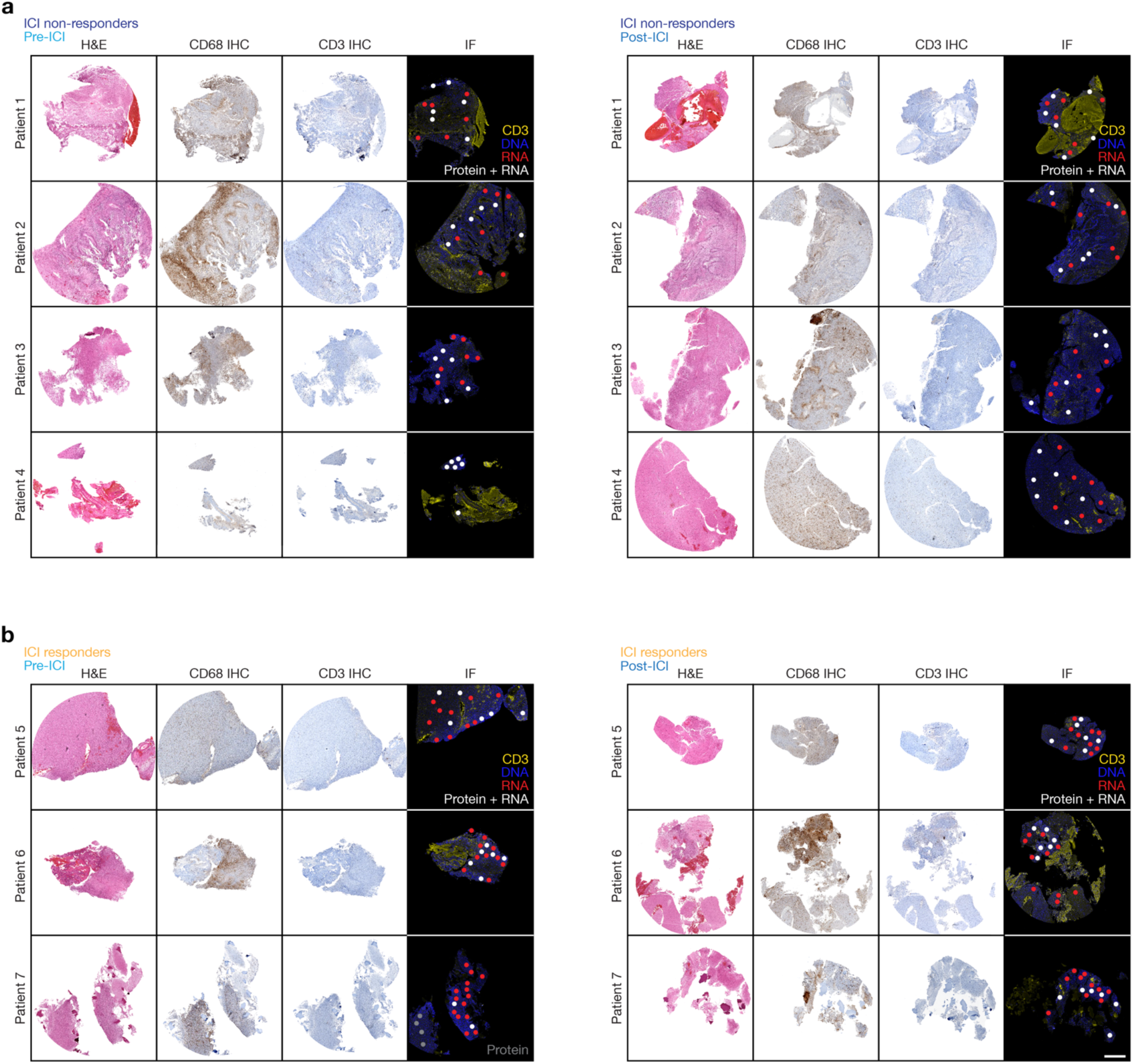
Human glioblastoma tissue sections pre-ICI and post-ICI. **a**, H&E, CD68 IHC, and CD3 IF sections for ICI responders pre-ICI (left) and post-ICI (right) showing region-of-interest (ROI) location for spatial proteomics (white circles) and spatial transcriptomics (white and red circles). **b**, H&E, CD68 IHC, and CD3 IF sections for ICI non-responders pre-ICI (left) and post-ICI (right) showing ROI location for spatial proteomic (white circles) and spatial transcriptomic (white and red circles) analyses. Scale bar, 1mm.

**Extended Data Fig. 2.**
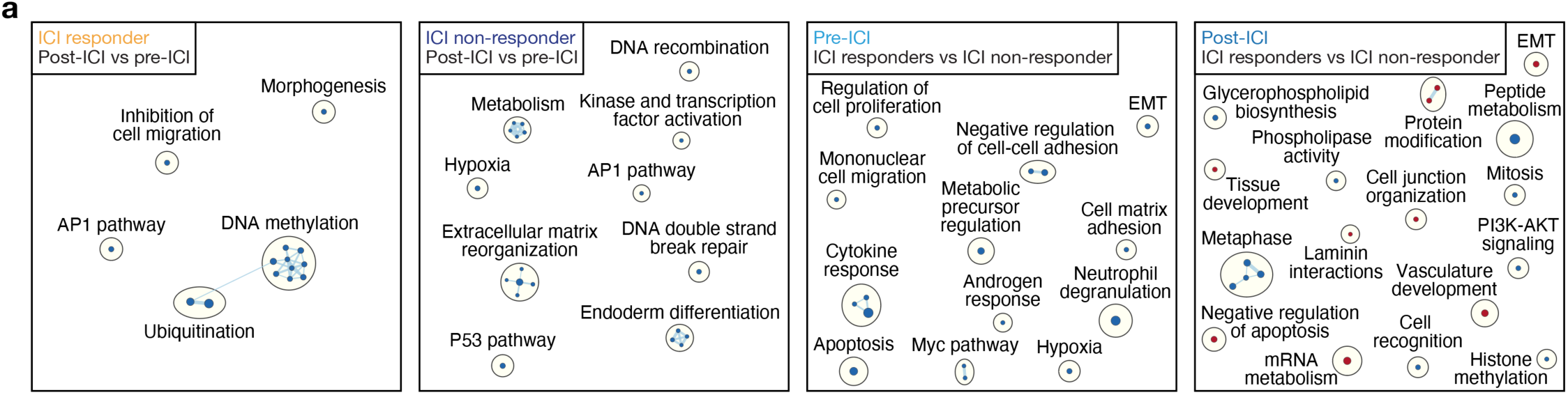
Gene circuit network changes between ICI responders and non-responders before and after ICI treatment. **a**, Network of gene circuits mapping differentially expressed genes between ICI responders (far left) and non-responders (middle left) post-ICI and pre-ICI specimens, and differentially expressed genes between pre-ICI (middle right) and post-ICI (far right) specimens from ICI responders and non-responders. Nodes represent pathways and edges represent shared genes between pathways (p≤0.05, FDR≤0.05). n = 14 match-paired glioblastoma samples from 7 patients.

**Extended Data Fig. 3.**
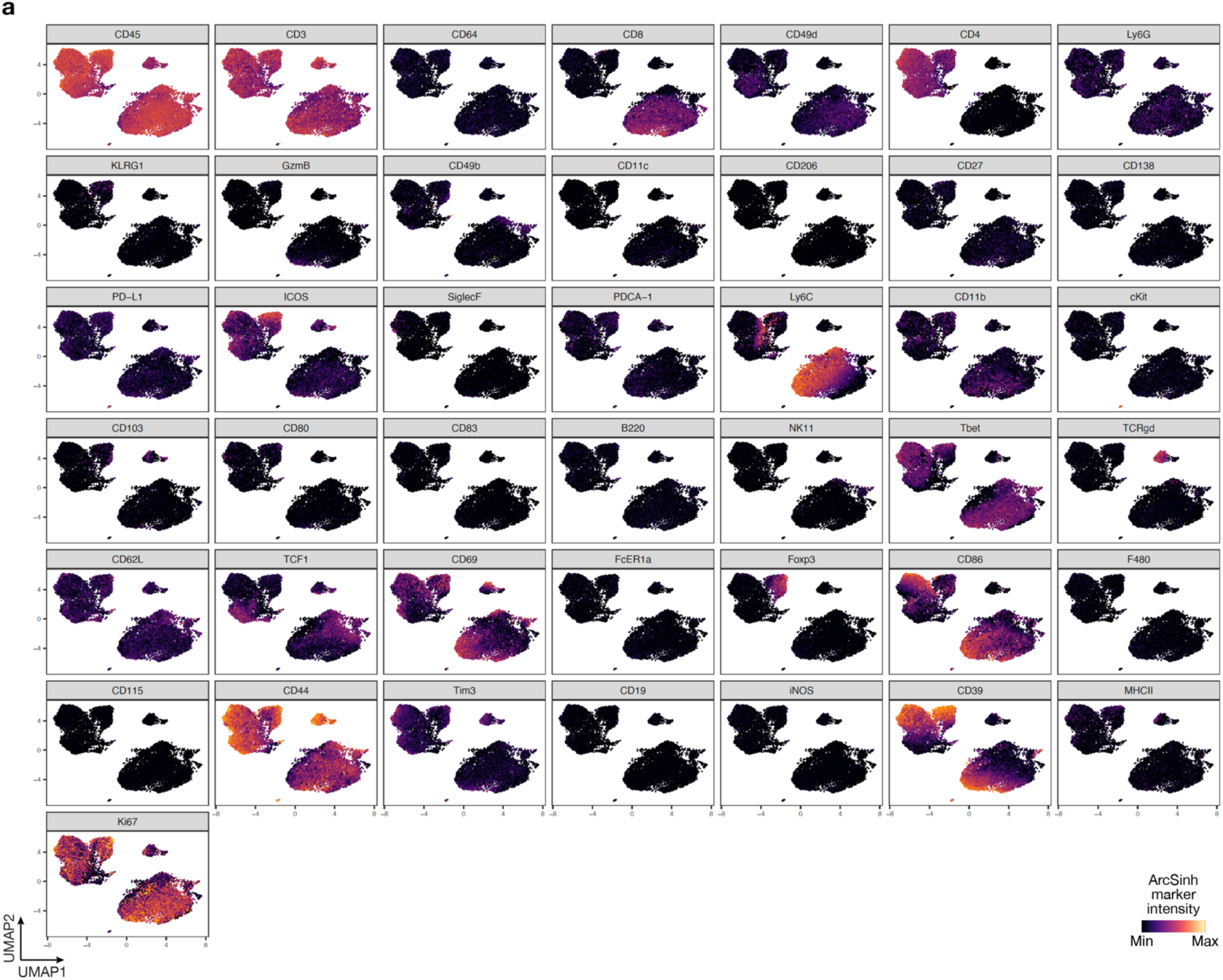
UMAP markers for CyTOF gating of T cells from intracranial mouse glioblastoma allografts. **a**, UMAP cell density plots showing protein expression used for assigning cell types in Fig. 4e.

**Extended Data Fig. 4.**
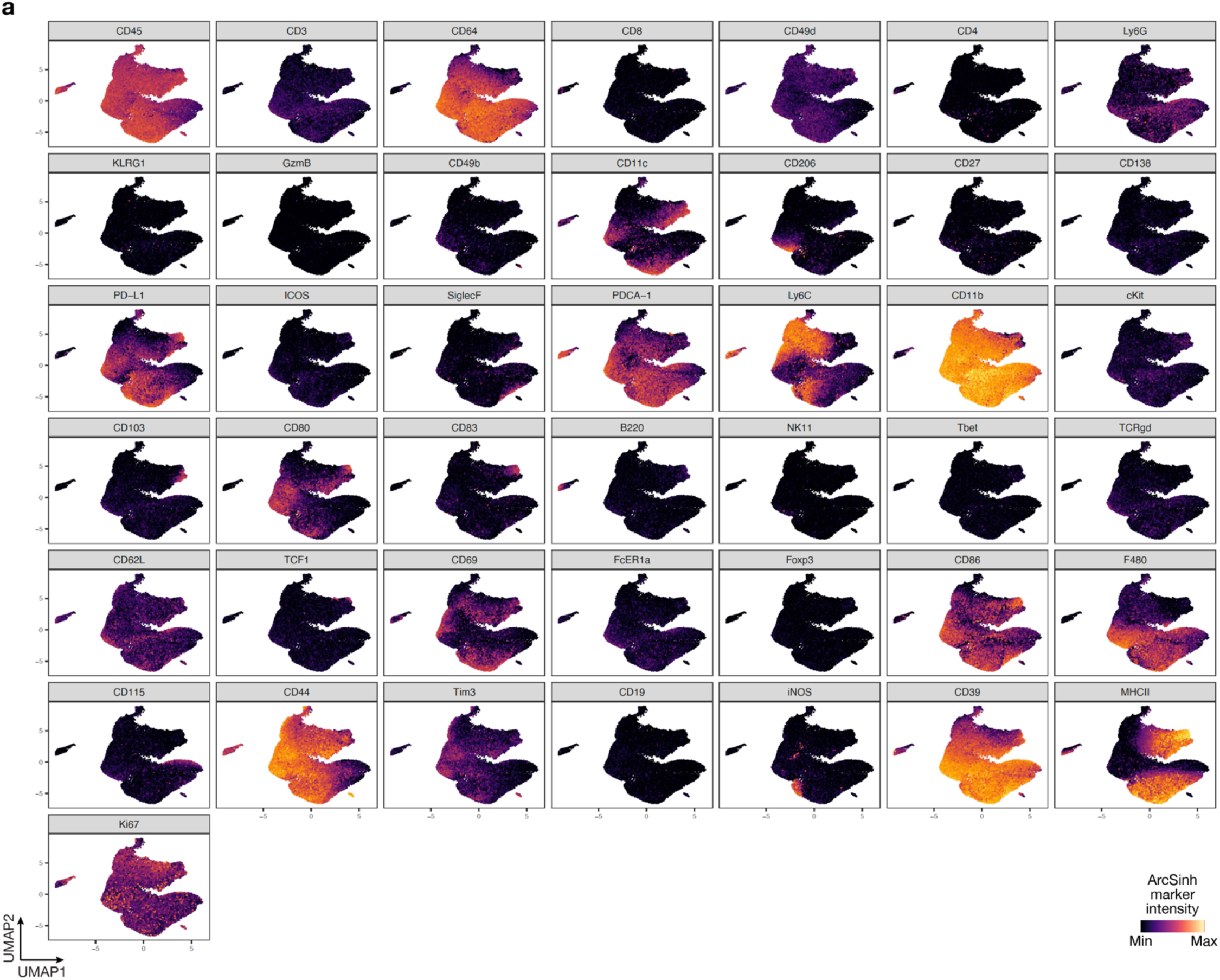
UMAP markers for CyTOF gating of myeloid cells from intracranial mouse glioblastoma allografts. **a**, UMAP cell density plots showing protein expression used for assigning cell types in Fig. 4e.

**Extended Data Fig. 5.**
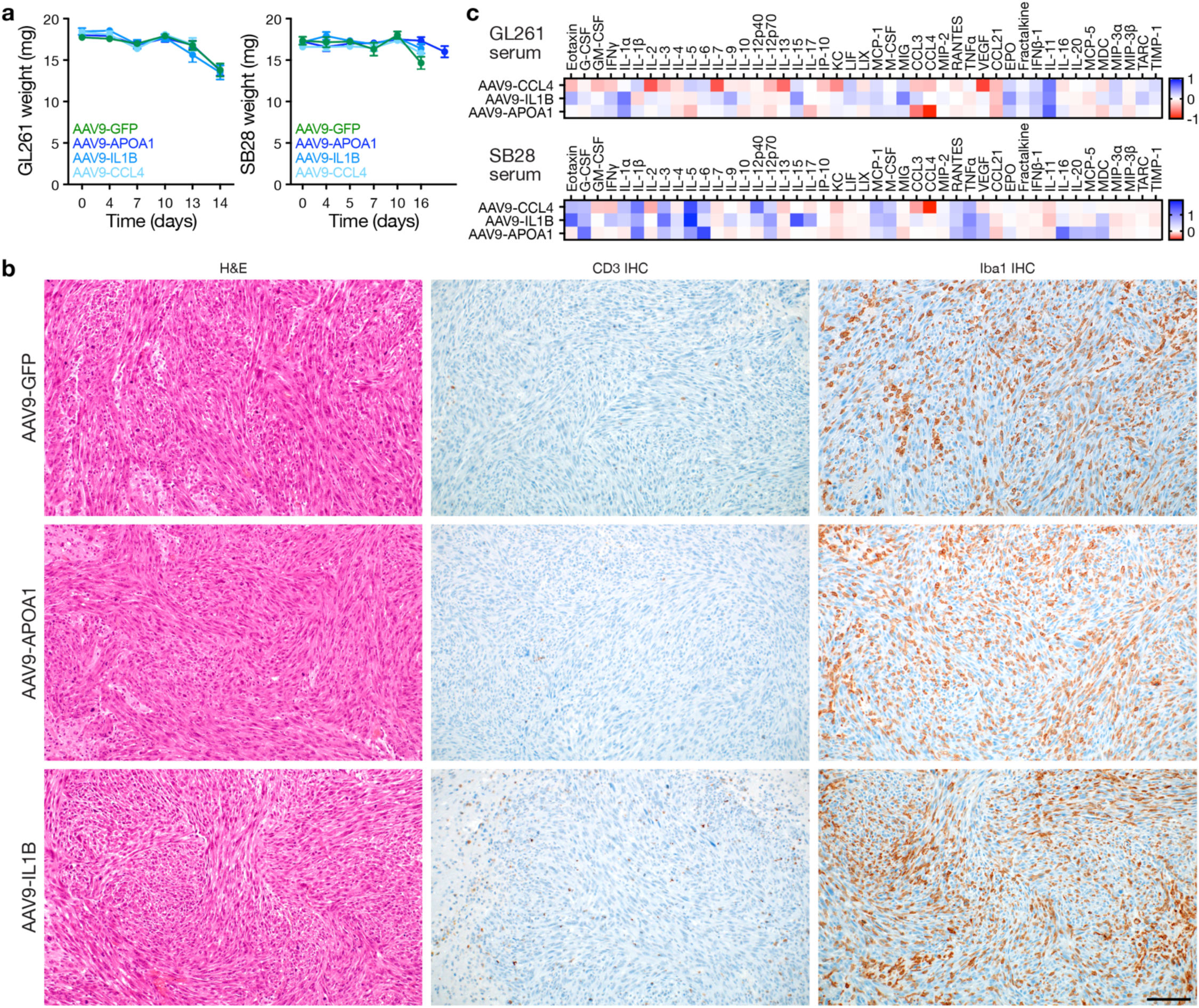
Intratumor cytokine gene therapy for intracranial glioblastoma allografts. **a**, Body weights for C57J/B6 WT mice harboring intracranial GL261 (left) or SB28 (right) intracranial allografts treated with intratumor CED cytokine gene therapy (n=10 mice/condition). See also Fig. 5c. **b**, H&E, CD3 IHC, and Iba1 IHC sections from SB28 allografts treated with intratumor AAV9 gene therapy showing modest CD3 infiltrate and robust Iba1 infiltrate in regions of viable tumor after AAV9-APOA1 or AAV9-IL1B compared to AAV9-GFP control (2x10^11^ viral genomes per mouse). Scale bar, 100µm. **c**, Heatmap of cytokine changes from multiplexed bead assays of sera from GL261 (top) or SB28 (bottom) intracranial mouse glioblastoma allografts after intratumor treatments with AAV9 gene therapies (n=3 mice/condition) vs AAV9-GFP (n=3 mice) showing a lack of consistent alterations in cytokine levels following intratumor gene therapy treatments. Legend shows mean log10 fold change with gene therapy treatments vs AAV9-GFP control.

**Extended Data Fig. 6.**
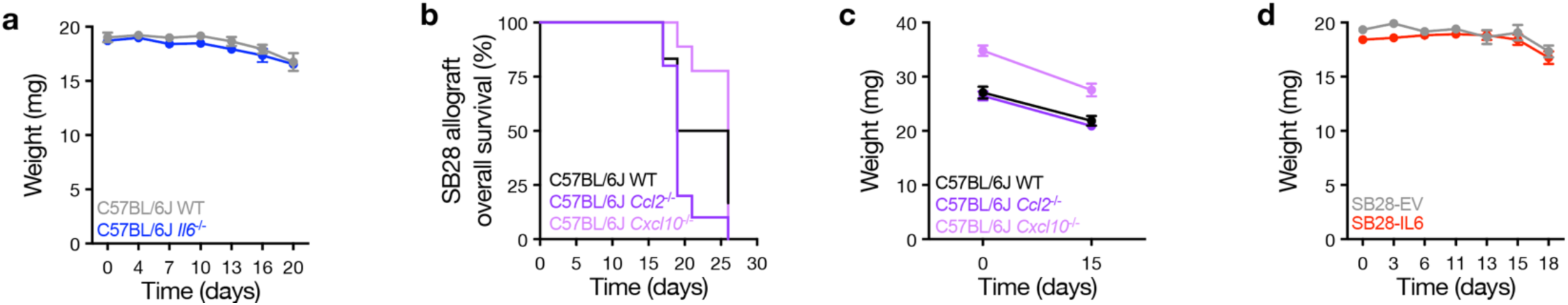
CCL2 and CXCL10 from the tumor microenvironment do not contribute to glioblastoma overall survival. **a**, Body weights for C57J/B6 WT mice (n=8) vs C57J/B6 *Il6^-/-^*mice (n=8) harboring intracranial SB28 glioblastoma allografts. See also Fig. 6g. **b**, Kaplan-Meier curves for overall survival from SB28 intracranial glioblastoma allografts in C57J/B6 WT (n=8 mice) vs C57J/B6 *Ccl2^-/-^* mice (n=8 mice) vs *Cxcl10^-/-^* mice (n=8 mice) showing no change in overall survival for mice lacking CCL2 or CXCL10 in the tumor microenvironment. Log rank test. **c**, Body weights for C57J/B6 WT (n=8 mice) vs C57J/B6 *Ccl2^-/-^* mice (n=8 mice) vs *Cxcl10^-/-^* mice (n=8 mice) harboring intracranial SB28 glioblastoma allografts. **d**, Body weights for C57J/B6 WT mice harboring SB28 intracranial glioblastoma allografts with doxycycline-induced *IL6* (n=10 mice) or EV (n=10 mice) overexpression. See also Fig. 6i.

**Extended Data Fig. 7.**
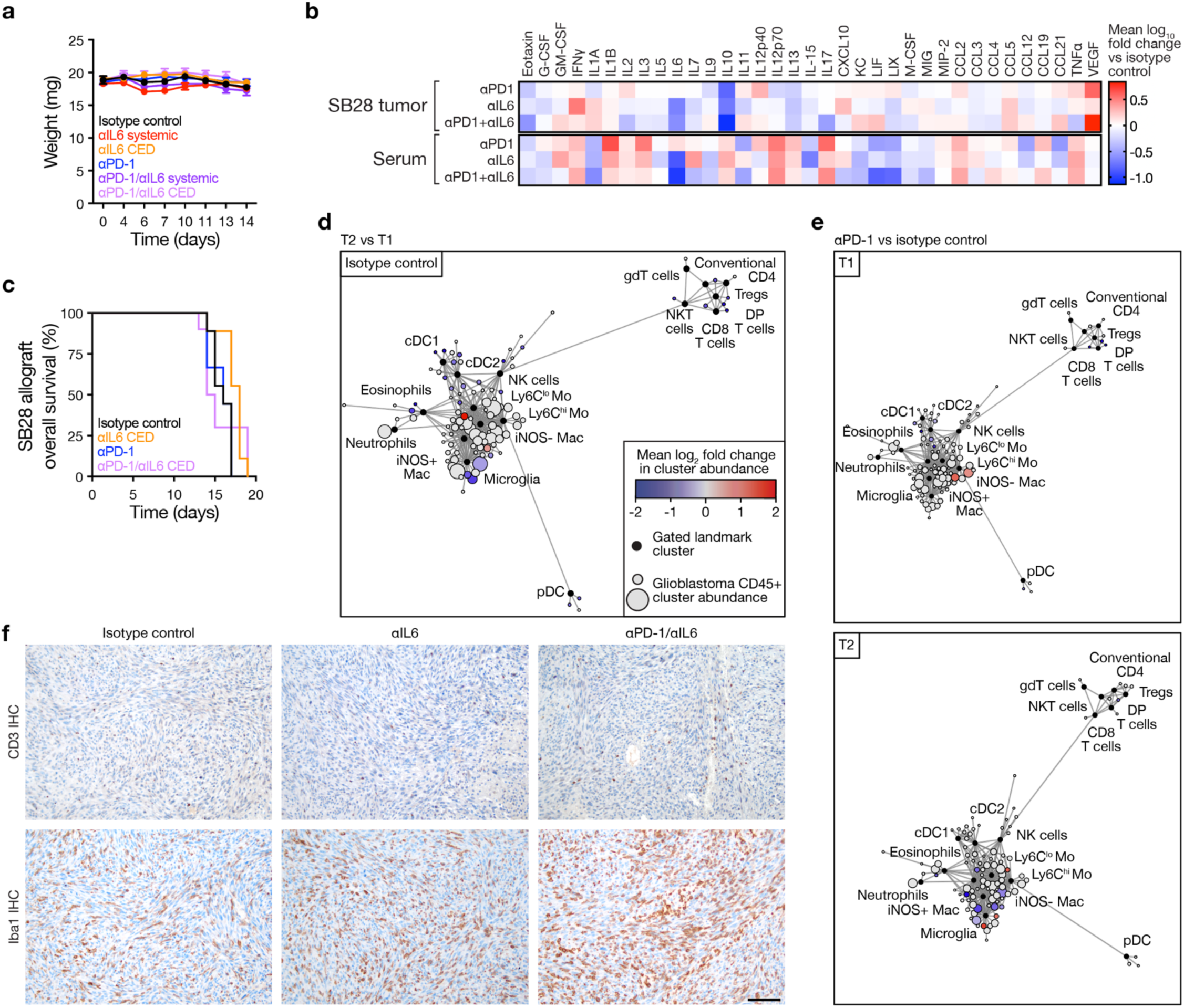
Intratumor IL6 blockade does not prolong glioblastoma overall survival and immune checkpoint inhibition in the absence of systemic IL6 blockade does not reprogram the glioblastoma immune microenvironment. **a**, Body weights for C57J/B6 WT mice (n=8-9 mice/condition) bearing SB28 intracranial mouse glioblastoma allografts treated with isotype control, systemic αIL6 monotherapy, systemic αPD-1 monotherapy, intratumor CED αIL6 monotherapy, or intratumor vs systemic αIL6 in combination systemic αPD-1 therapy. See also Fig. 7b. **b**, Heatmap of cytokine changes from multiplexed bead assays of SB28 intracranial glioblastoma allograft tumor lysates (top) or sera (bottom) after systemic monotherapy or combined antibody treatments (n=8 mice/condition) showing suppression of IL10 with all antibody treatments and suppression of IL6 with αIL6 treatments. **c**, Kaplan-Meier curves for overall survival from SB28 intracranial glioblastoma allografts showing no difference in overall survival after intratumor CED treatment with αIL6 (156.75 μg in 15μL) either alone or in combination with systemic αPD-1 therapy (250 μg biweekly i.p.) compared to isotype control treatments (n=8-9 mice/condition). **d**, CD45+ immune cell CyTOF scaffold plot (n=104,388 cells) from SB28 intracranial glioblastoma allografts treated with isotype control at T2 vs T1 timepoints from Fig. 7b showing microglia in the tumor immune microenvironment decrease over time. **e**, CD45+ immune cell CyTOF scaffold plots from SB28 intracranial glioblastoma allografts treated with αPD-1 monotherapy vs isotype control at T1 (top, n=117,751 cells) or T2 (bottom, n= 194,205 cells) timepoints from Fig. 7b showing minimal change in the tumor immune microenvironment with αPD-1 monotherapy. **f**, CD3 IHC and Iba1 IHC sections from intracranial glioblastoma SB28 allografts in C57BL/6J mice treated with isotype control, systemic αIL6 monotherapy, or combined αPD-1/αIL6 therapy showing increases in Iba1 macrophages and CD3+ T cells following combination therapy. Scale bar, 100µm.

**Extended Data Fig. 8.**
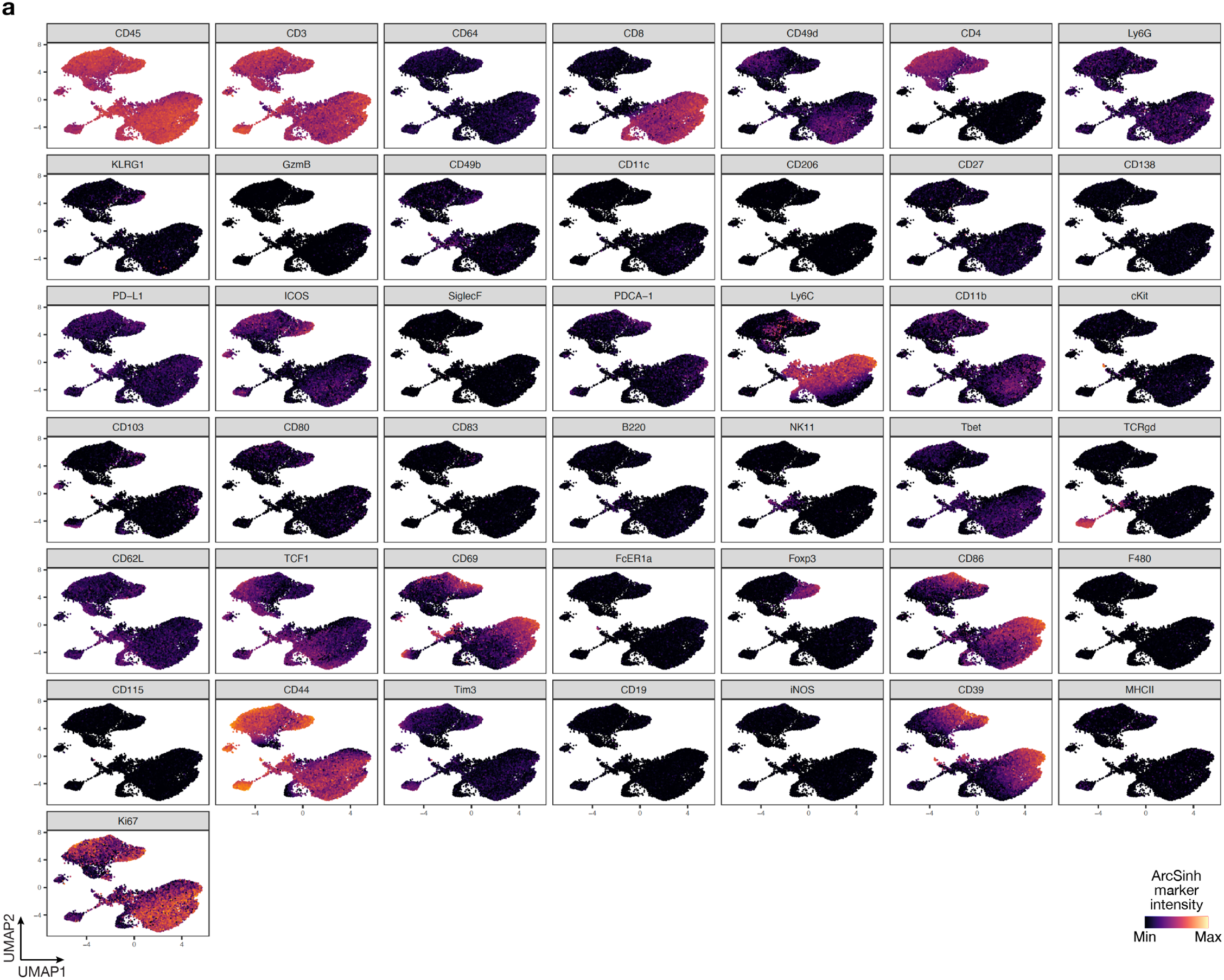
UMAP markers for CyTOF gating of T cells from intracranial mouse glioblastoma allografts after IL6 blockade and immune checkpoint inhibition. **a**, UMAP cell density plots showing protein expression used for assigning cell types in Fig. 7d.

**Extended Data Fig. 9.**
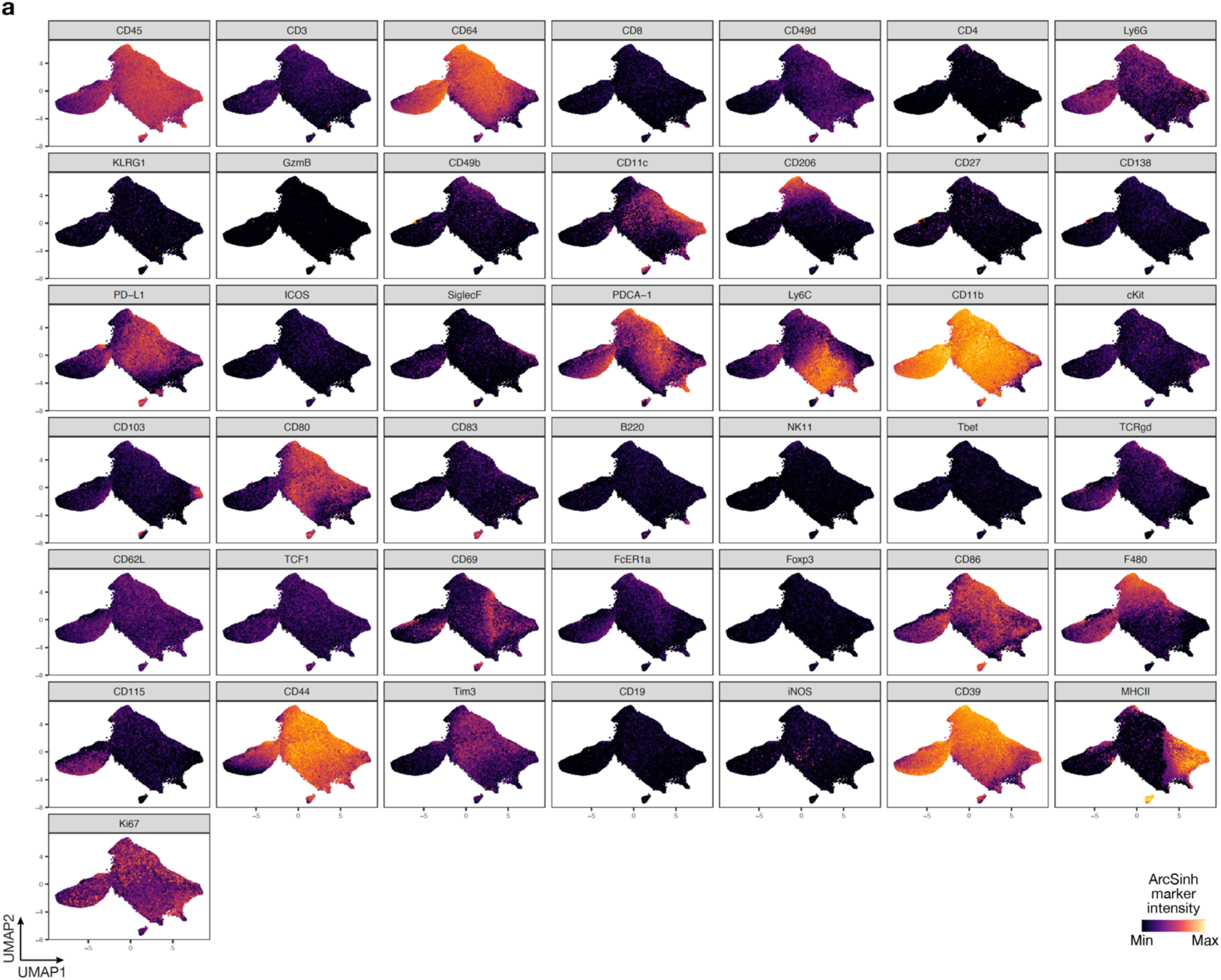
UMAP markers for CyTOF gating of myeloid cells from intracranial mouse glioblastoma allografts after IL6 blockade and immune checkpoint inhibition. **a**, UMAP cell density plots showing protein expression used for assigning cell types in Fig. 7d.

**Extended Data Fig. 10.**
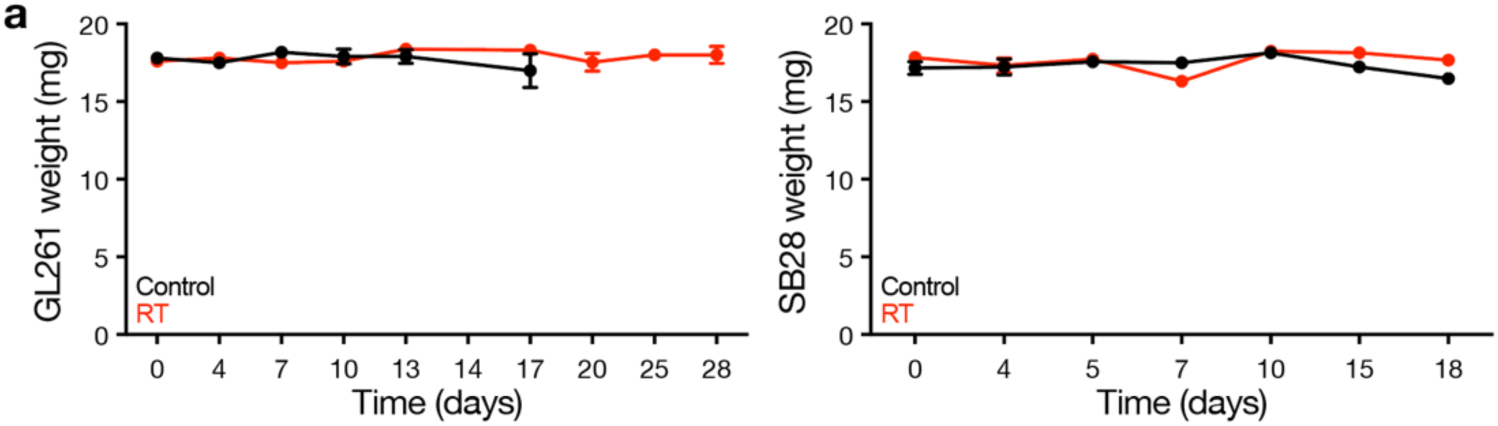
Body weights for immunocompetent mice treated with radiotherapy. **a**, Body weights for C57J/B6 WT mice harboring intracranial GL261 (left) or SB28 (right) intracranial allografts treated with radiotherapy (RT, 18Gy/1Fx) vs no-treatment control (n=10 mice/condition). See also Fig. 7a.

**Extended Data Fig. 11.**
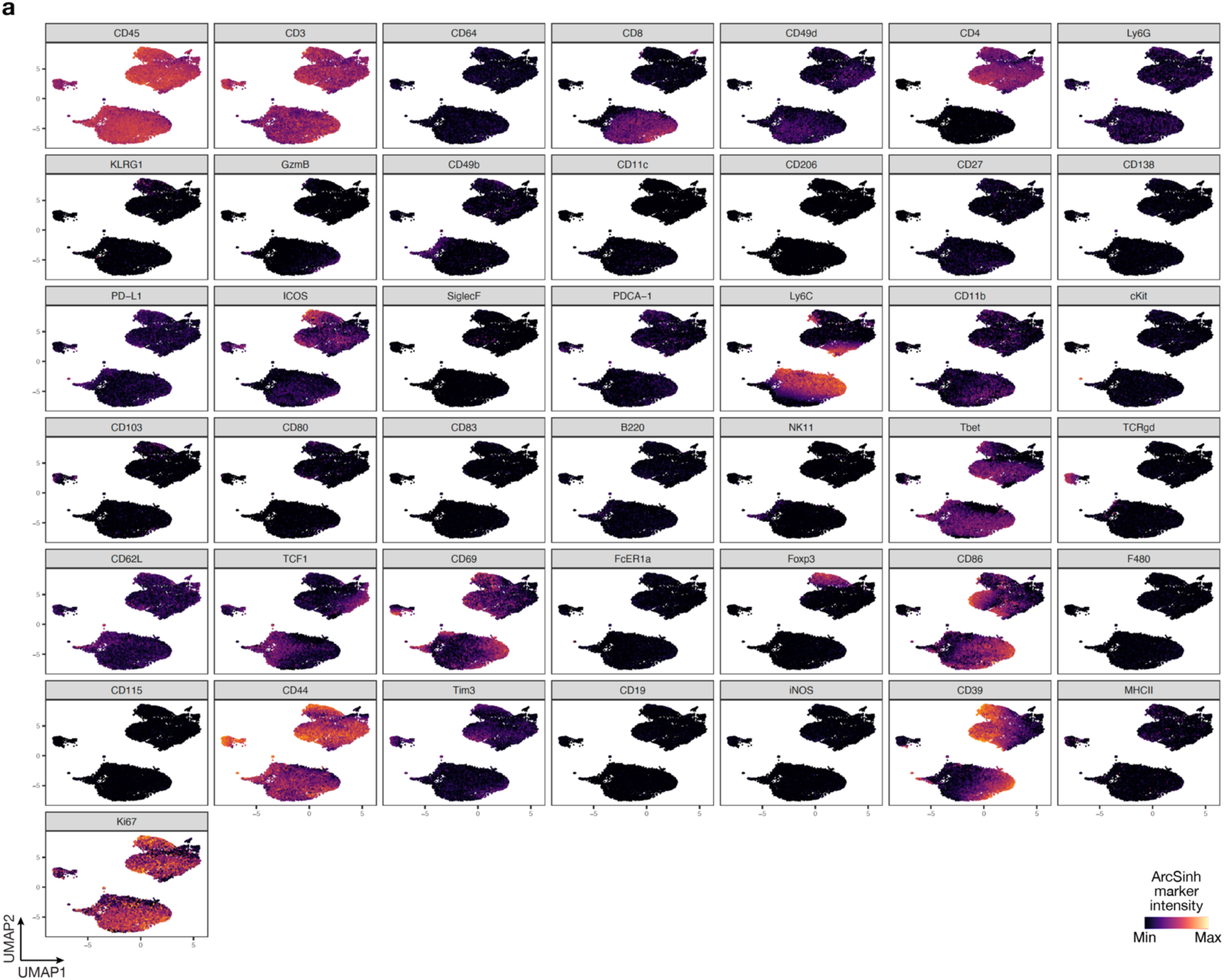
UMAP markers for CyTOF gating of T cells from intracranial mouse glioblastoma allografts after radiotherapy, IL6 blockade, and immune checkpoint inhibition. **a**, UMAP cell density plots showing protein expression used for assigning cell types in Fig. 8c.

**Extended Data Fig. 12.**
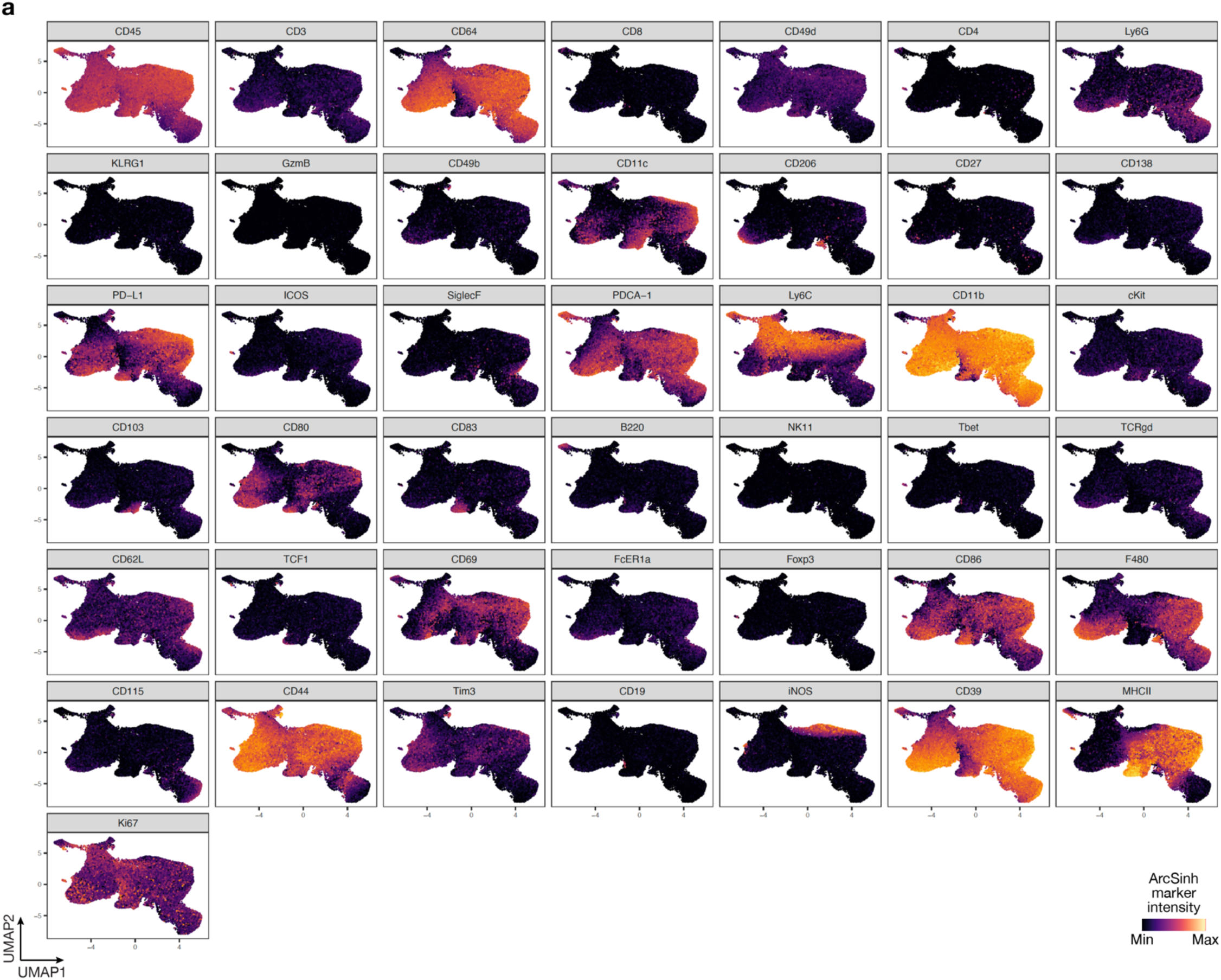
UMAP markers for CyTOF gating of myeloid cells from intracranial mouse glioblastoma allografts after radiotherapy, IL6 blockade, and immune checkpoint inhibition. **a**, UMAP cell density plots showing protein expression used for assigning cell types in Fig. 8c.

**Extended Data Fig. 13.**
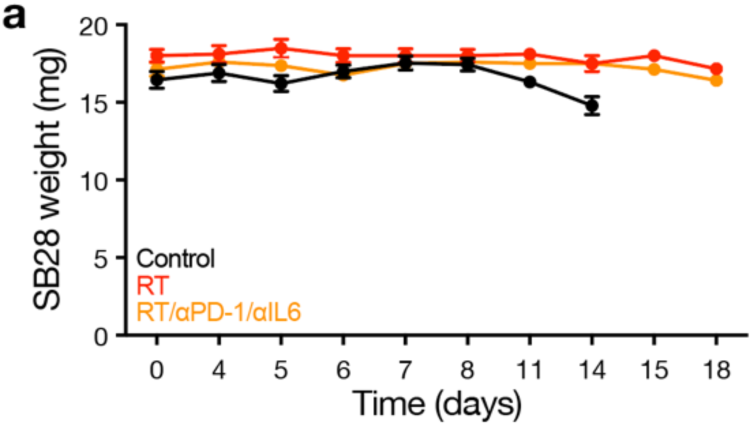
Body weights for immunocompetent mice treated with radiotherapy, IL6 blockade, and immune checkpoint inhibition. **a**, Body weights for C57J/B6 WT mice harboring intracranial SB28 glioblastoma allografts treated with radiotherapy (RT, 18Gy/1Fx) vs RT plus αPD-1/αIL6 therapy (250 μg biweekly i.p. for up to four weeks for each antibody) vs no-treatment control (n=8-10 mice/condition). See also Fig. 8g.

